# Genome-wide analysis of the polyamine oxidase gene family in wheat (*Triticum aestivum* L.) reveals involvement in temperature stress response

**DOI:** 10.1101/2020.07.06.189209

**Authors:** Fatemeh Gholizadeh, Ghader Mirzaghaderi

## Abstract

Amine oxidases (AOs) including copper containing amine oxidases (CuAOs) and FAD-dependent polyamine oxidases (PAOs) are associated with polyamine catabolism in the peroxisome, apoplast and cytoplasm and play an essential role in growth and developmental processes and response to biotic and abiotic stresses. Here, we identified *PAO* genes in common wheat (*Triticum aestivum*), *T. urartu* and *Aegilops tauschii* and reported the genome organization, evolutionary features and expression profiles of the wheat PAO genes (*TaPAO*). Expression analysis using publicly available RNASeq data showed that *TaPAO* genes are expressed redundantly in various tissues and developmental stages. A large percentage of *TaPAOs* respond significantly to abiotic stresses, especially temperature (i.e. heat and cold stress). Some *TaPAOs* were also involved in response to other stresses such as powdery mildew, stripe rust and *Fusarium* infection. Overall, *TaPAOs* may have various functions in stress tolerances responses, and play vital roles in different tissues and developmental stages. Our results provided a reference for further functional investigation of TaPAO proteins.

## Introduction

Common wheat (*Triticum aestivum* L., 2*n* = 6*x* = 42; AABBDD genome), is one of the most important cereal crops. It is constantly exposed to abiotic and biotic stresses such as heat, cold, salinity, drought and various fungal diseases. These stresses reduce growth and yield and may cause plant death. Therefore it is essential to understand how wheat adapts and survives in stressful environments, and to develop methods to increase its tolerance under environmental stresses [1].

Polyamines (PAs), are small aliphatic amines of low molecular weight that are involved in various developmental processes in living organisms. Main PAs in cells include diamine putrescine (Put), triamine spermidine (Spd), tetramines spermine (Spm), cadaverine (Cad) and thermospermine (T-Spm). Due to their cationic nature, polyamines are capable of binding to negatively charged molecules such as RNA and DNA and affect gene expression, protein synthesis and regulation of ion channels [2]. De novo production of PAs in plants includes Put production directly from ornithine by ornithine decarboxylase (ODC), or indirectly from arginine by arginine decarboxylase (ADC) [1]. Put is then converted into Spd by spermidine synthase with the addition of an amino propyl moiety donated by decarboxylated S-adenosyl methionine (dcSAM). Similarly, Spm (and its isomer T-Spm) is formed from Spd via Spm synthase, with the same amino propyl group rendered by dcSAM [3, 4] (Fig. 1).

**Fig. 1.**
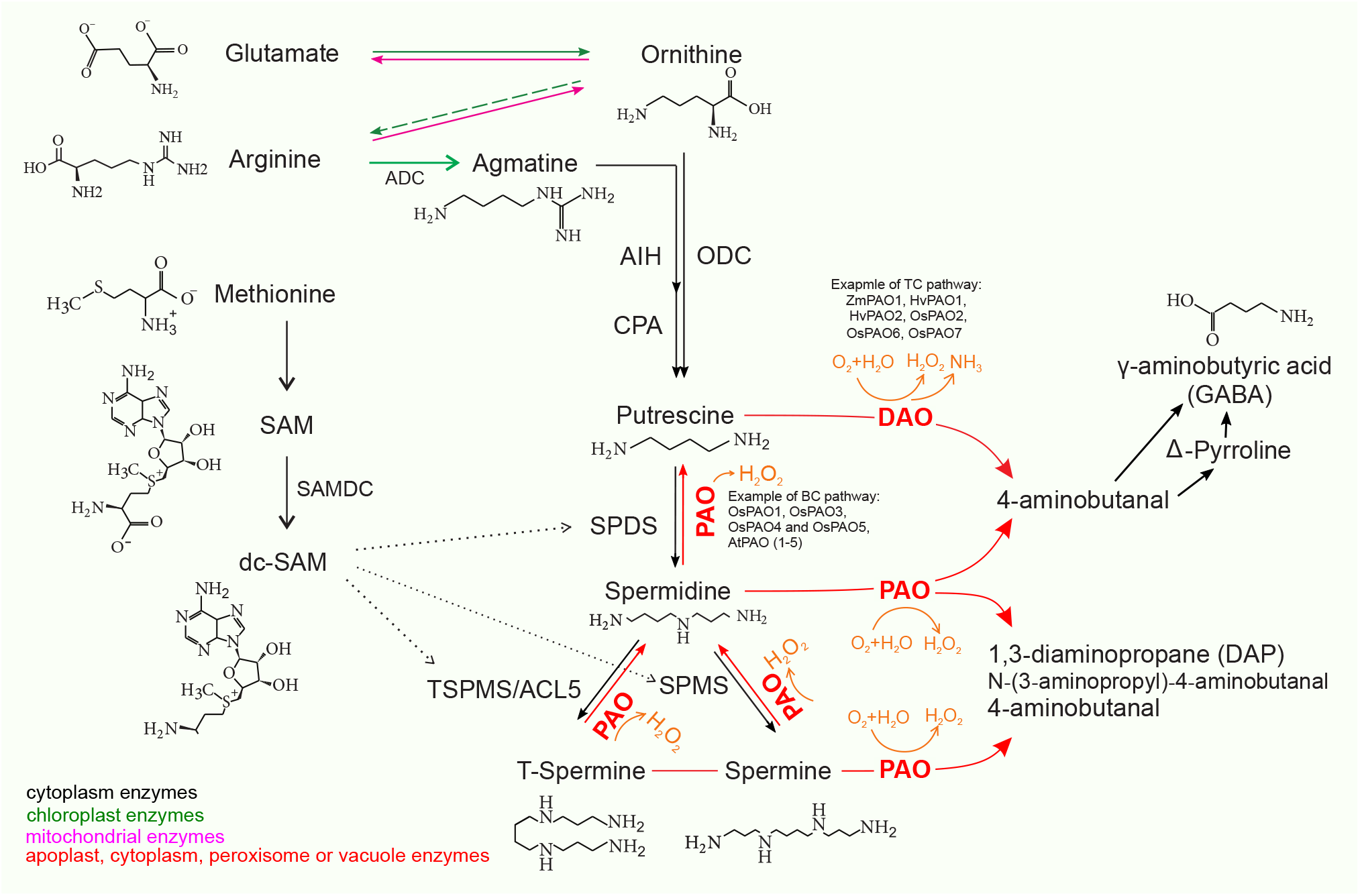
Polyamine biosynthesis in plants. ADC, arginine decarboxylase; AIH, agmatine iminohydrolase; CPA, N-carbamoyl putrescine amidohydrolase; dcSAM: decarboxylated S-adenosylmethionine; SAM: S-adenosylmethionine; SAMDC: S-adenosylmethionine decarboxylase; SPDS: spermidine synthase; SPMS: spermine synthase; TSPMS: thermospermine synthase; spermidine synthase: SPDS; spermine synthase: SPMS; PAO: polyamine oxidase. The donor of the aminopropyl groups is dc-SAM, which is formed by decarboxylation of SAM, through an enzymatic reaction catalyzed by SAMDC. The aminopropyltransferases donating aminopropyl residue to Put or Spd for production of Spd or Spm are SPDS and SPMS.

PAs can be oxidized by copper-containing diamine oxidases (CuAOs or DAOs) and flavin-containing (FAD-containing) polyamine oxidases (PAOs) [5]. DAOs mainly catalyze the oxidation of Put and Cad producing 4-aminobutanal, ammonia (NH_3_) and hydrogen peroxide (H_2_O_2_) [6, 7]. POAs are divided into two major groups. The first group catalyzes Spd and Spm to produce 1,3-diaminopropane (DAP), H_2_O_2_, and N-(3-aminopropyl)-4-aminobutanal or 4-aminobutanal, which is referred to as the terminal catabolism (TC) pathway [5, 7, 8]. The second group is involved in the back conversion (BC) pathway by converting Spm back to Spd and Spd to Put [7, 9].

Plants accumulate osmolyte compounds in response to abiotic stresses such as drought and salinity. Major cellular osmolytes including proline, glycine betaine, and PAs are found in plants, animals, and bacteria [10]. In plants, PAs are essential for development and stress response. Many plant processes such as embryogenesis, organogenesis, particularly flower initiation and development, fruit setting and ripening, as well as leaf senescence, require PAs [3, 11]. Cells need to maintain the homeostasis of PAs through their modulation, biosynthesis, conjugation, and transport, since high concentrations of polyamines are highly toxic [12].

Spd and Spm and Put levels are differentially regulated by environmental stresses [13], although the mechanism of PA action in response to stresses still remain unclear. Put levels are increased with low potassium (K^+^) availability in plants suggest that Put and its catabolites possess a potential in controlling cellular K^+^ and Ca^2+^ [14]. During drought, the PA pathway is activated which leads to a Put to Spd canalization that is ABA-dependent. Drought tolerant and sensitive cultivars seem to be different in their capacity to accumulate different PAs over a minimum threshold [15].

H_2_O_2_ produced through PA oxidation is involved in a hyper-sensitive (HR) reaction that can lead to bacterial pathogen tolerance [16]. Exogenous Spm results in HR-mediated resistance of *Arabidopsis* leaves to cucumber mosaic virus via the induction of the expression of some H_2_O_2_-dependent signaling components and transcription factors. Addition of a PAO inhibitor represses the activation of defense genes and alleviates ROS generation and HR, confirming that PAO is involved in the resistance response [17]. There is evidence that PA oxidation in the apoplast together with the generated reactive oxygen species (ROS) are involved in programmed cell death (PCD) and xylem differentiation [3]. The transcript levels of PA synthesis genes, and the activities of corresponding enzymes are responsive to stresses, providing a relationship between polyamine and stresses [1]. Plant PAOs play significant roles in metal (e.g. aluminum, copper, and cadmium) toxicity tolerance [18–22]. In wheat, the cell wall-bound PAO (CW-PAO) oxidized Spd and generated H_2_O_2_ under aluminum toxicity but Put application resulted in plant tolerance against Aluminum-induced oxidative stress via inhibiting PAO activity and hence lowering H_2_O_2_ production [20].

*PAO* genes have been isolated and characterized from several model plants. One of the first polyamine oxidases identified was a FAD-based PAO in maize apoplast, a 53-kDa monomeric glycoprotein enzyme [23]. Most of the identified plant *PAO* genes such as *A. thaliana AtPAO1* to *AtPAO5* are involved in the BC pathway. AtPAO1 and AtPAO5 are located in the cytoplasm, while AtPAO2, AtPAO3 and AtPAO4 have a peroxisomal localization [24–26]. AtPAO1 is involved in biotic and abiotic stress tolerance and may play roles in root development and fertility. On the other hand AtPAO2 might be involved in root, shoot, leaf, and flower development. AtPAO3 and AtPAO4 are expressed in all tissues and whole growth stages and show similar expression patterns [27, 28]. Rice harbors seven *PAO* genes. *OsPAO3* and *OsPAO5* are very similar and highly expressed in both the seedling stage and in mature plants, while the other *PAO* members are only expressed at very low levels in all plant tissues. OsPAO4 and OsPAO5 prefer to use Spm and T-Spm as substrates, but cannot oxidize Spd to Put. Therefore, OsPAO3 catalyzes a full BC-type pathway, while OsPAO4 and OsPAO5 only catalyze a partial BC-type pathway [4].

In the present study, polyamine oxidase genes were identified in *T. aestivum*, *T. urartu* and *Aegilops tauschii* using bioinformatic approaches and their gene structure, conserved protein motifs and domains and phylogenetic relationships were analyzed. Furthermore, we examined the expression of the wheat *PAO* genes over different tissues and developmental stages and in response to biotic and abiotic stresses.

## Materials and Methods

### Identification of *PAO* genes

Polyamine oxidase genes of common wheat (*T. aestivum*) and its relatives *T. urartu* and *Ae. tauschii*, were identified by BLASTP search, Hidden Markov Model (HMM) analysis and validation of conservative domains. For this, the *Arabidopsis* and rice PAO protein sequences (supplementary File 1) were used as queries to perform BLASTP searches against the *T. aestivum*, *T. urartu* and *Ae. tauschii* genome (E-value < 1e-5) in the EnsemblPlants database at https://plants.ensembl.org. Furthermore, an HMM matrix of five AtPAO and seven OsPAO protein sequences was used to search the PAO proteins in jackhmmer (https://www.ebi.ac.uk/tools/hmmer/search/jackhmmer) [29]. We then selected the unique sequences of the above two search results and checked them for the presence of each of the amine oxidase domains (Pfam: PF01593) alone or in combination with copper amine oxidase (N2 and/or N3-terminal), using the Pfam (https://pfam.xfam.org) and InterPro (http://www.ebi.ac.uk/interpro) databases. Proteins with amine oxidase in combination with other extra domains were excluded, as such architectures are known to have functions different from PAO. For example, plant lysine histone demethylases which possess an additional SWIRM domain are involved in demethylation of mono- and di-methylated lysines of histones [30]. Other described genes such as zeta-carotene desaturase, protoporphyrinogen oxidase, prolycopene isomerase and protein FLOWERING locus D-like protein were also excluded.

### Identification of orthologs and homoeologs

*PAO* homoeologous genes and pairwise gene orthologs among *T. aestivum*, *T. urartu*, *Ae. tauschii*, *A. thaliana* and *Oryza sativa* were identified through the “homoeologous” and “orthologoues” links in the gene-based display of the EnsemblPlants summary page for each target gene. PAO genes were mapped to their respective locus in the wheat genome in a circular diagram using shinyCircos [31] where homoeologous chromosomes were aligned close together and banded according to the general FISH patterns of p*Ta*535-1 and (GAA)_10_ probes.

### Characterization of *TaPAO* genes

Characteristics of each of the identified amino oxidase proteins such as isoelectric point (pI), amino acid sequence length (AA) and molecular weight (MW) were obtained from the ProtParam website at https://web.expasy.org/protparam [32]. A GFF3 annotation file containing the locations of *TaPAOs* in genome and their structural information was extracted from the wheat GFF3 file and the exon-intron structures was displayed using the Gene Structure Display Server (GSDS, http://gsds.cbi.pku.edu.cn [33]. The conserved domains of the TaPAO protein sequences were searched from Pfam [34] and MEME [35] websites and the resulting files were visualized in TBtools software [36]. Wheat and rice PAO protein sequences were also aligned in Jalview [37] and the locations of the domains identified by MEME, were determined on the alignment output file.

### Phylogenetic analysis

Multiple sequence alignment of the full-length protein sequences of the identified PAO proteins was performed using the “msa” package [38] of R version 3.6.1 (The R Project for Statistical Computing, Vienna, Austria). Subsequently, a neighbor-joining tree was obtained with 100 bootstrap replicates using the “ape” package [39] and used to generate a tree in R using the “ggtree” package [40].

### Expression analysis of *TaPAO* genes using RNAseq

RNAseq data of 30 *TaPAO* genes was retrieved from www.wheat-expression.com [41] as processed expression values in transcripts per million (TPM) for all the available tissues and developmental stages [42] and for response to different stresses including *Fusarium* [43, 44], cold [45], *Zymoseptoria* [46], heat and drought [47], phosphorous starvation [48], powdery mildew [49] and PEG (https://www.ebi.ac.uk/ena/browser/view/PRJNA306536). *TaPAO* gene expression values were transformed and used to generate barplots in R. Count matrix data of all experiments were also downloaded and used for differential gene expression analysis, using the DESeq2 package [50] to statistically compare the mean expression level of each *TaPAO* gene between control and stress conditions. A heatmap was generated from log_2_(TPM+1) transformed values of *TaPAO* genes over developmental stages using R package “pheatmap”. Ternary plots were generated from the stress response data using the R package ggtern [51]. For this, genes with zero expression in all homoeologs were excluded.

### Detecting alternative splicing events among *TaPAOs*

Wheat genome sequences and annotations (IWGSC RefSeq v1.0) [52] were downloaded from https://plants.ensembl.org/info/website/ftp/index.html. In order to detect and visualize the alternative splice variants, we firstly downloaded RNAseq reads [SRP043554, 45] from https://www.ebi.ac.uk. RNAseq data belong to the wheat plants (‘Manitou’ cultivar) in three-leaf stage at normal (grown at 23°C for 4 weeks after germination) and cold stress (grown at 23°C for 2 weeks followed by 4°C for another 2 weeks) conditions. After removing the low quality reads and inspecting for adapter sequences, the raw RNA sequence data from each sample were mapped to the wheat reference genome using HISAT2 and transcripts were assembled and merged using StringTie with default settings [53]. Normalization of abundance estimates as FPKM (fragments per kilobase of transcript per million mapped reads) values, differential gene and transcript expression analysis and graphical displaying of alternative splice variants were done using the “ballgown” package [54].

## Results

### Identification of PAO proteins in common wheat, *T. urartu* and *Ae. tauschii*

BLASTP and the Hidden Markov Model (HMM) matrix of *Arabidopsis* and rice polyamine oxidase genes (Supplementary File 1) was used to search the amino oxidase proteins in common wheat, *Ae. tauschii* and *T. urartu* protein databases. In total, after verification of the identified sequences for the presence of each amino_oxidase domain (Pfam: PF01593) or copper amine oxidase-catalytic domain, either alone or in combination with copper amine oxidase (N2 and/or N3-terminal), 30 *PAO* genes in *T. aestivum*, 6 *PAO* genes in *T. urartu* and 8 *PAO* genes in *Ae. tauschii* were identified. These genes were named *TaPAO1* to *TaPAO11*, followed by the name of the harbouring chromosome. For those identified *PAO* genes which were orthologous to rice *PAOs*, the same numbers were assigned as for the rice *PAO* genes (Table 1).

**Table 1.** Information and physicochemical characteristics of *PAO* genes in bread wheat, *T. urartu* and *Ae. tauschii*. Notes: AA, amino acid sequence length; MW, molecular weight; pI, isoelectric point. ASN: alternative splice variants. “1” indicates only a single transcript. *: wheat *PAO* genes that are confidently orthologous with the corresponding rice *PAOs*. **ASN:** alternative splice variants from EnsemblPlants. **D:** gene direction, ‘+’: forward. ‘−’: reverse. **ASN**:** alternative splice variants identified in ‘Manitou’ cultivar from experiment SRP043554.

| Species | Name | Transcript ID | AA | MW (kDa) | pI | ASN | D | ASN** |
| --- | --- | --- | --- | --- | --- | --- | --- | --- |
| *T. urartu* | TuPAO5 | TRIUR3_11268-T1 | 520 | 57376.68 | 5.34 | 1 | + | - |
|  | TuPAO9 | TRIUR3_14057-T1 | 504 | 56367.99 | 6.45 | 1 | + | - |
|  | TuPAO6 | TRIUR3_12020-T1 | 454 | 50979.07 | 7.12 | 1 | + | - |
|  | TuPAO3 | TRIUR3_18876-T1 | 484 | 53632.17 | 5.34 | 1 | - | - |
|  | TuPAO4 | TRIUR3_11269-T1 | 520 | 57376.68 | 5.34 | 1 | + | - |
|  | TuPAO10 | TRIUR3_14834-T1 | 490 | 55178.33 | 5.36 | 1 | + | - |
| *Ae. tauschii* | AetPAO4-2D | AET2Gv21199400.1 | 490 | 53358.12 | 5.36 | 10 | + | - |
|  | AetPAO3-2D | AET2Gv21031900.5 | 513 | 57023.23 | 5.51 | 36 | - | - |
|  | AetPAO5-2D | AET2Gv21199100.12 | 492 | 54373.33 | 5.51 | 15 | + | - |
|  | AetPAO9-4D | AET4Gv20654900.7 | 526 | 59025.88 | 6.55 | 12 | - | - |
|  | AetPAO6-7D | AET7Gv21301800.1 | 498 | 55934.48 | 5.99 | 6 | - | - |
|  | AetPAO11-7D | AET7Gv20928100.8 | 503 | 56533.91 | 5.64 | 11 | - | - |
|  | AetPAO1-3D | AET3Gv20612000.2 | 517 | 55184.35 | 5.09 | 3 | + | - |
|  | AetPAO10-4D | AET4Gv20866800.1 | 245 | 26562.27 | 5.79 | 10 | + | - |
| *T. aestivum* | TaPAO8-1A | TraesCS1A02G407600.1 | 585 | 61964.34 | 7.93 | 1 | + | 3 |
|  | TaPAO8-5B | TraesCS5B02G529400.1 | 585 | 62050.42 | 8.40 | 1 | - | 4 |
|  | TaPAO8-5D | TraesCS5D02G528500.1 | 582 | 61688.97 | 7.59 | 1 | - | 2 |
|  | TaPAO10-4B | TraesCS4B02G385300.1 | 481 | 53676.73 | 5.76 | 1 | - | 1 |
|  | TaPAO10-5A | TraesCS5A02G549600.1 | 495 | 55509.74 | 5.60 | 1 | + | 1 |
|  | TaPAO11-7A | TraesCS7A02G378800.1 | 457 | 51891.59 | 5.55 | 1 | - | 1 |
|  | TaPAO11-7B | TraesCS7B02G280700.1 | 477 | 54210.02 | 5.55 | 1 | - | 2 |
|  | TaPAO11-7D | TraesCS7D02G375700.1 | 503 | 56533.91 | 5.64 | 2 | - | 2 |
|  | TaPAO2-2A | TraesCS2A02G053400.1 | 340 | 37651.87 | 5.02 | 1 | - | 1 |
|  | TaPAO3-2A* | TraesCS2A02G467300.1 | 484 | 53646.20 | 5.34 | 1 | - | 2 |
|  | TaPAO3-2B* | TraesCS2B02G490100.1 | 484 | 53604.16 | 5.34 | 1 | - | 3 |
|  | TaPAO3-2D* | TraesCS2D02G467300.1 | 484 | 53632.17 | 5.34 | 1 | - | 2 |
|  | TaPAO4-2A* | TraesCS2A02G548200.1 | 490 | 53266.07 | 5.37 | 1 | + | 2 |
|  | TaPAO4-2B* | TraesCS2B02G579100.1 | 490 | 53312.05 | 5.35 | 1 | + | 1 |
|  | TaPAO4-2D* | TraesCS2D02G549300.1 | 540 | 58748.37 | 5.64 | 1 | + | 2 |
|  | TaPAO5-2A* | TraesCS2A02G548100.1 | 487 | 53768.67 | 5.44 | 1 | + | 4 |
|  | TaPAO5-2B* | TraesCS2B02G579000.1 | 526 | 57604.88 | 5.45 | 1 | + | 2 |
|  | TaPAO5-2D* | TraesCS2D02G549200.1 | 492 | 54373.33 | 5.51 | 3 | + | 4 |
|  | TaPAO6-7A* | TraesCS7A02G539200.1 | 508 | 56928.63 | 6.58 | 2 | - | 2 |
|  | TaPAO6-7B* | TraesCS7B02G461800.1 | 495 | 55486.12 | 6.40 | 1 | + | 1 |
|  | TaPAO6-7D* | TraesCS7D02G524900.1 | 498 | 55946.45 | 5.99 | 1 | + | 1 |
|  | TaPAO9-2A | TraesCS2A02G159500.1 | 474 | 49845.89 | 6.27 | 1 | + | 1 |
|  | TaPAO9-2B | TraesCS2B02G185100.1 | 471 | 49635.65 | 6.11 | 1 | + | 1 |
|  | TaPAO1-3A* | TraesCS3A02G250700.1 | 510 | 54902.17 | 5.49 | 1 | + | 1 |
|  | TaPAO1-3B* | TraesCS3B02G280200.1 | 507 | 54518.62 | 5.22 | 1 | + | 1 |
|  | TaPAO1-3D* | TraesCS3D02G251100.1 | 491 | 52509.23 | 5.07 | 1 | + | 1 |
|  | TaPAO7-4A | TraesCS4A02G039600.1 | 468 | 52554.66 | 9.30 | 3 | + | 3 |
|  | TaPAO7-4B | TraesCS4B02G265900.1 | 493 | 55334.93 | 7.23 | 1 | - | 1 |
|  | TaPAO7-4D | TraesCS4D02G265800.1 | 493 | 55354.78 | 6.52 | 1 | - | 1 |
|  | TaPAOUn | TraesCSU02G062000.1 | 585 | 61995.31 | 7.58 | 1 | + | 1 |

### Phylogeny and characterization of *PAO* genes

The sequence length of TaPAO proteins ranged from 340 (TaPAO2-2A) to 585 (TaPAO8-1A, TaPAO8-5B and TaPAOUn) amino acids. The average molecular weight was 54.68 kDa, varying between 37.87 kDa (TaPAO2-2A) and 62.42 kDa (TaPAO8-5B). The isoelectric points (pI) of TaPAO members ranged from 5.02 (TaPAO2-2A) to 9.30 (TaPAO7-4A), with an average of 6.11, showing a weak acidity (Table 1). In order to identify the evolutionary relationships between PAO members, a phylogenetic tree of 56 PAO protein sequences belonging to *T. aestivum, T*. *urartu, A*. *tauschii, O. sativa* and *A. thaliana* was constructed using protein sequences based on the neighbor-joining method. The tree clustered the PAOs into seven clades (Fig. 2). Clade I contains four TaPAO11 homoeologs plus AetPAO11-7D of *Ae. tauschii*. Clade II was composed of TaPAO9 and TaPAO8 homoeologs, TaPAOUn, and TaPAO2-2A. clade III was composed of TaPAO1 homoeologs plus AetPAO1-3D of *Ae. tauschii* together with OsPAO1 and AtPAO5. Clade IV contained TaPAO4 and five homoeologs together with their orthologs from *T. urartu*, *Ae. tauschii* and *O. sativa* plus AtPAO4. Clade-V had eight members including TaPAO3 homoeologs together with their orthologs from *T. urartu*, *Ae. tauschii* and *O. sativa* plus AtPAO2 and 3. Clade VI contained only AtPAO1, which appeared significantly different from other characterized PAOs. Clade VII was the biggest clade with 17 PAO proteins including TAPAO6 and 7 homoeologs together with their *Ae. tauschii*, *T. urartu* orthologs. *O. sativa* OsPAO2, OsPAO6 and OsPAO7 proteins are also in the clade VII which are involved in the TC catabolism pathway (Fig. 1). Taken together, it seems that the identified wheat PAOs in the present study were not equally distributed among the different clades. Based on the retrieved data from EnsemblPlants, *TaPAO5-2D*, *TaPAO6-7A*, *TaPAO7-4A*, *TaPAO11-7D* and all the *Ae. tauschii* genes produces multiple splice variant (Table 1).

**Fig. 2.**
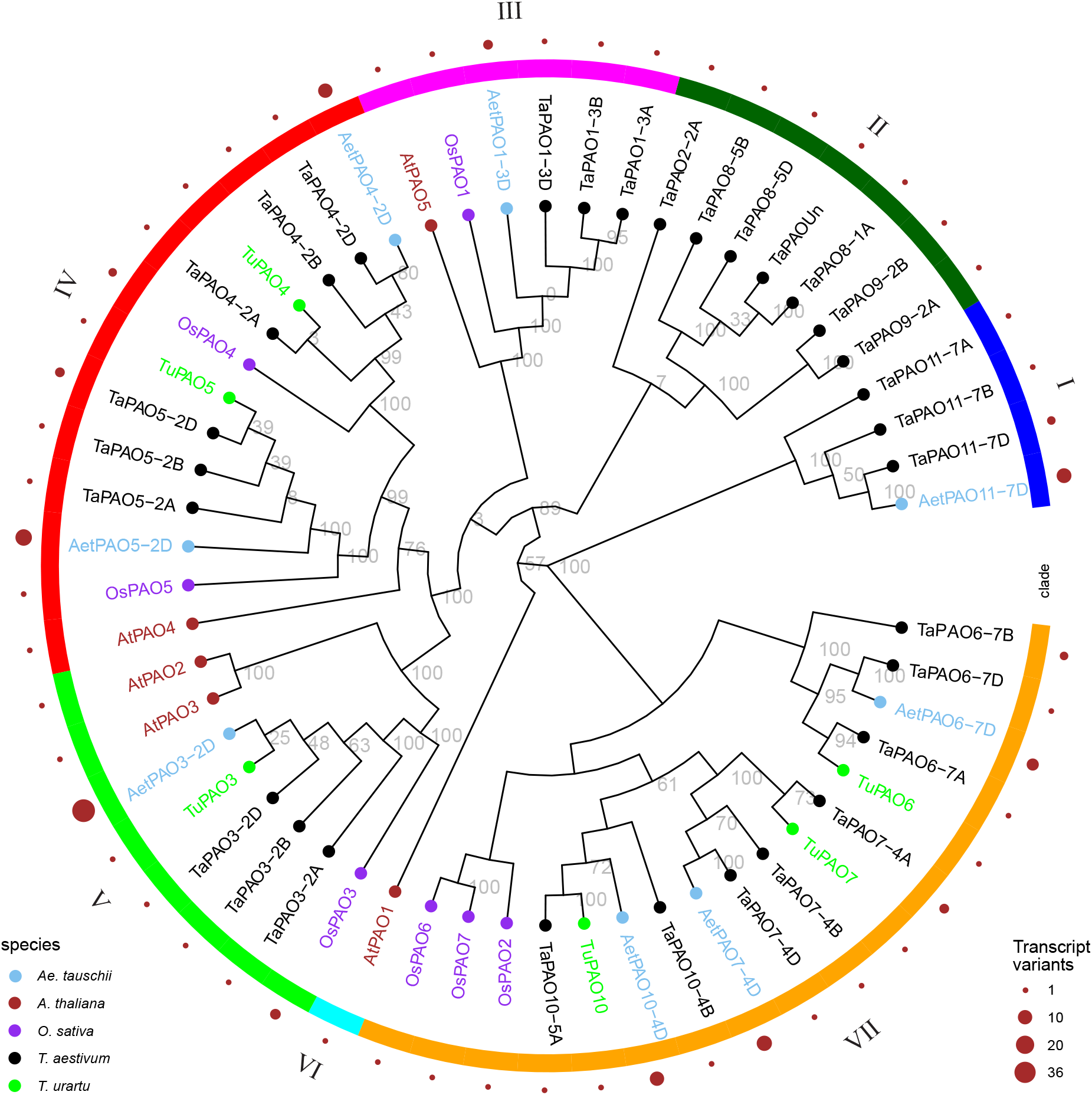
Phylogenetic tree of PAO proteins from *T. aestivum*, *T. urartu* and *Ae. tauschii*, *O. sativa* and *A. thaliana*.

### Analysis of chromosomal locations of *TaPAO* genes

A physical map of the location of the *TaPAO* genes on the A, B, and D chromosomes is illustrated in Fig. 3. The *TaPAO* genes were mapped to 16 wheat chromosomes plus the unassembled (Un) part of the genome. Homoeologs were connected using central links. Homoeologous chromosomes were aligned close together and banded according to the general FISH patterns of p*Ta*535-1 and (GAA)_10_ probes. The *TaPAO* genes showed uneven distribution across the A, B, and D subgenomes with a higher density on homoeologous group 2, and absence on chromosomes 1B, 1D and 6A, 6B and 6D. *TaPAO3*, *TaPAO4* and *TaPAO5* showed a similar exon/intron structure (Fig. 4) and were located together on the distal end of the long arm of homoeologous group 2, with the same order. *TaPAO6* and *TaPAO11* were also located close together on homoeologous group 7A, 7B and 7D but did not show noticeable structural similarity.

**Fig. 3.**
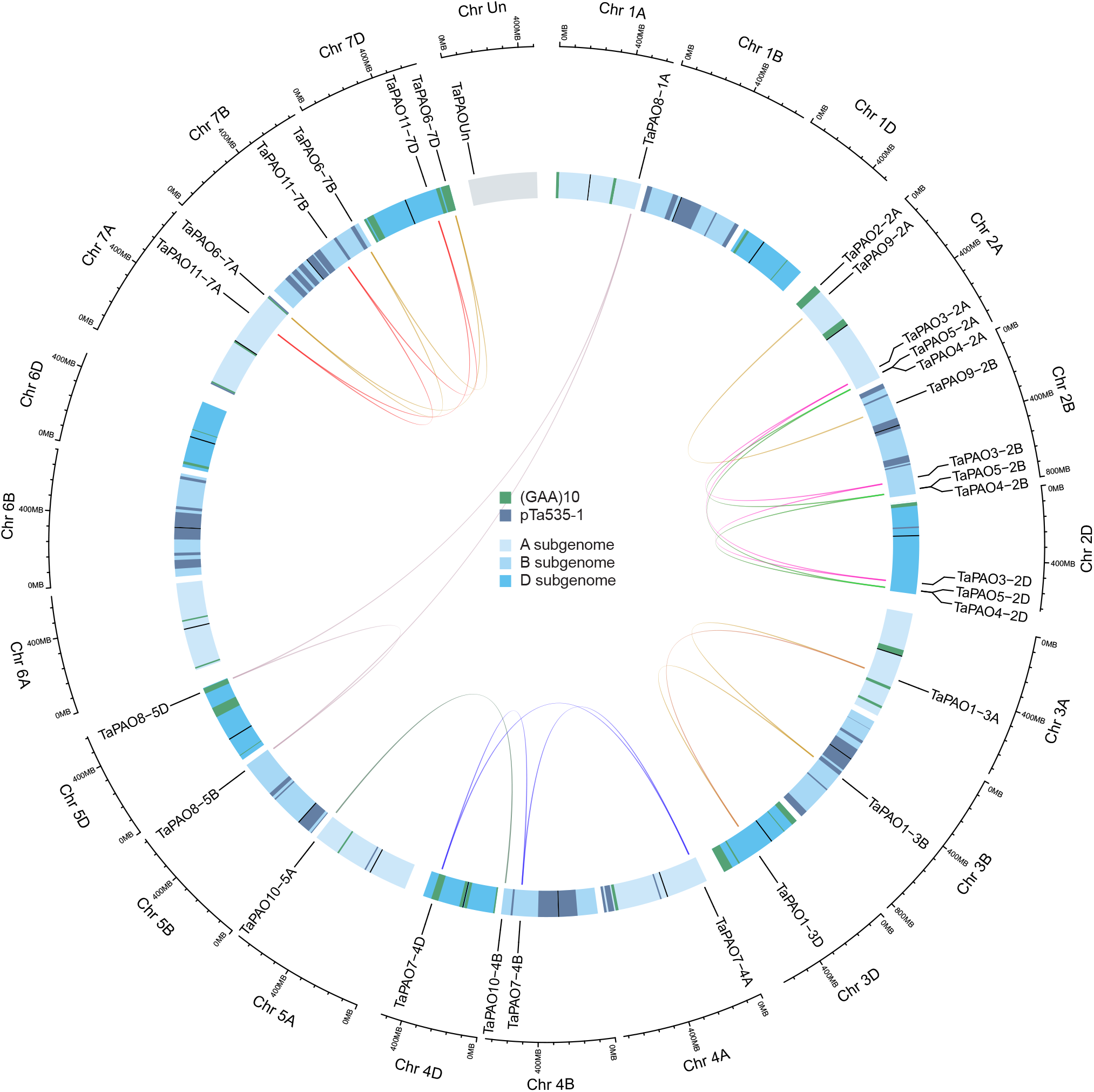
Chromosomal location of PAO genes on wheat chromosomes. Homoeologous genes were mapped to 16 wheat chromosomes (composed of A, B, and D subgenomes) plus one unassembled chromosome (Un) using shinyCircos. Homoeologs were connected using central links. Chromosome were banded according to p*Ta*535-1 (red bands) and (GAA)_10_ (blue bands) FISH patterns. Chromosome number is indicated outside the outer circle.

**Fig. 4.**
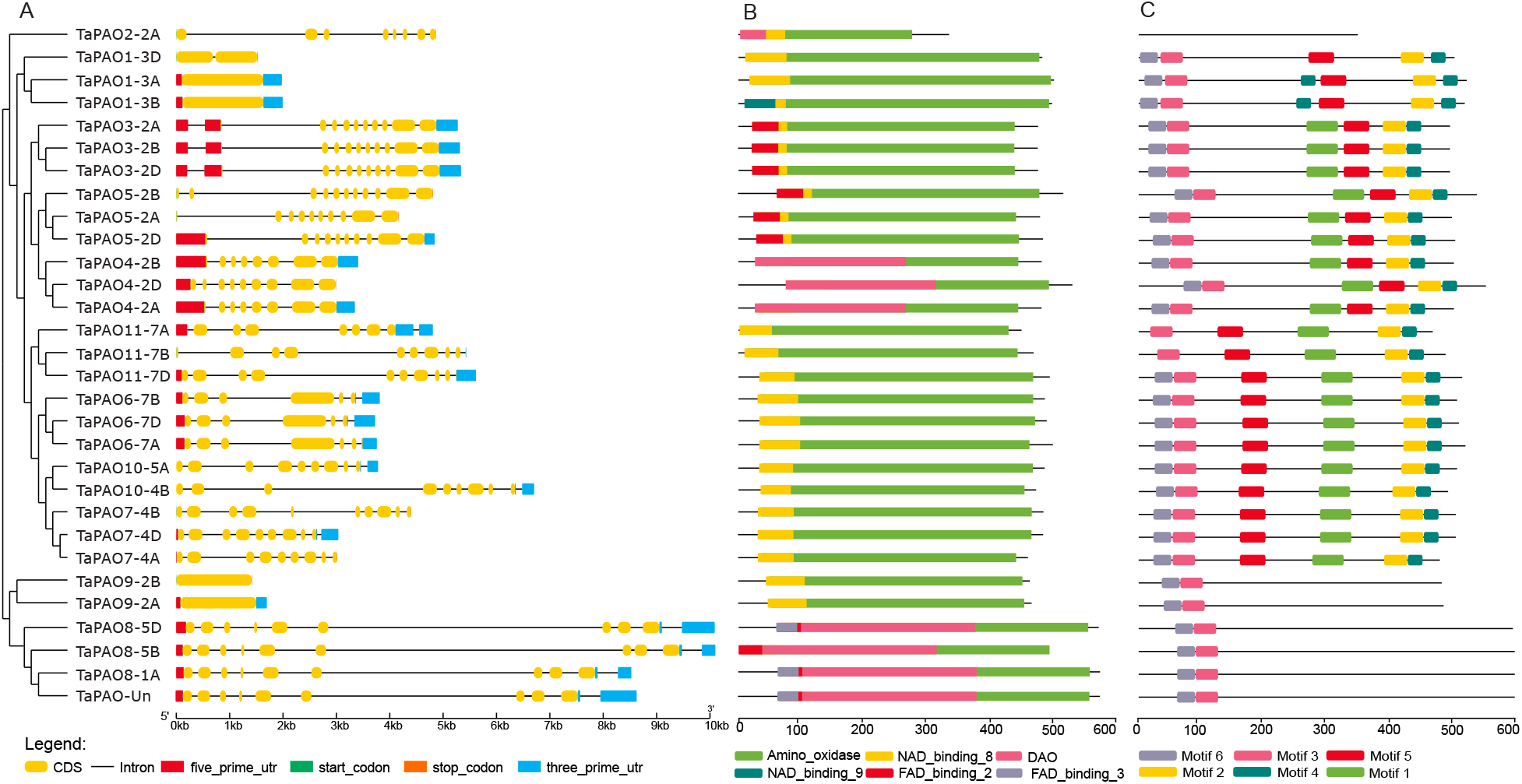
Gene structure, protein domain and motif analysis of *TaPAOs*. A) Exon–intron structures of *TaPAO* genes. B) Distribution of conserved domains within TaPAO proteins. C) Distribution of all motifs identified by MEME.

### Structure, domain and motif analysis of *TaPAO* genes

Exon–intron structural diversity within a gene family is an important clue for the evolutionary and functional analyses of gene family members. Gene structure, exons and introns were obtained for the identified 30 *TaPAO* genes to interrogate their genomic organization (Fig. 4A). Based on the wheat genome annotation, most *TaPAO* genes have introns in their structure and the number of exons varied from 1 (*TaPAO9-2A*, *TaPAO9-2B*, *TaPAO1-3A* and *TaPAO1-3B*) to 11 (*TaPAO5-2B*).

Protein domain analysis showed that most TaPAO members contained a typical amino_oxidase catalytic domain (alone or in combination with DAO) plus an NAD/FAD binding domain, with only TaPAO4-2A/-2B/-2D lacking an NAD/FAD binding domain (Fig. 4B). The MEME motif search tool identified six conserved motifs in TaPAO proteins (Fig. 5). The distribution patterns of these motifs in TaPAO proteins is shown in Fig. 4C. Motif 3 is present in all TaPAO proteins except TaPAO2-2A. Motif 6 uniformly distributed to all TaPAOs except TaPAO11-7A/-7B and TaPAO2-2A. Motif 1 was available in all TaPAO except TaPAO2-2A, TaPAO1-3A/3B/3D, TaPAO9-2A/2B, TaPAO8-1A/5B/5D and TaPAO-Un. Motifs 2, 4 and 5 were present in all TaPAOs except TaPAO2-2A, TaPAO9-2A/2B, TaPAO8-1A/5B/5D and TaPAO-Un (Fig. 4C).

**Fig. 5.**
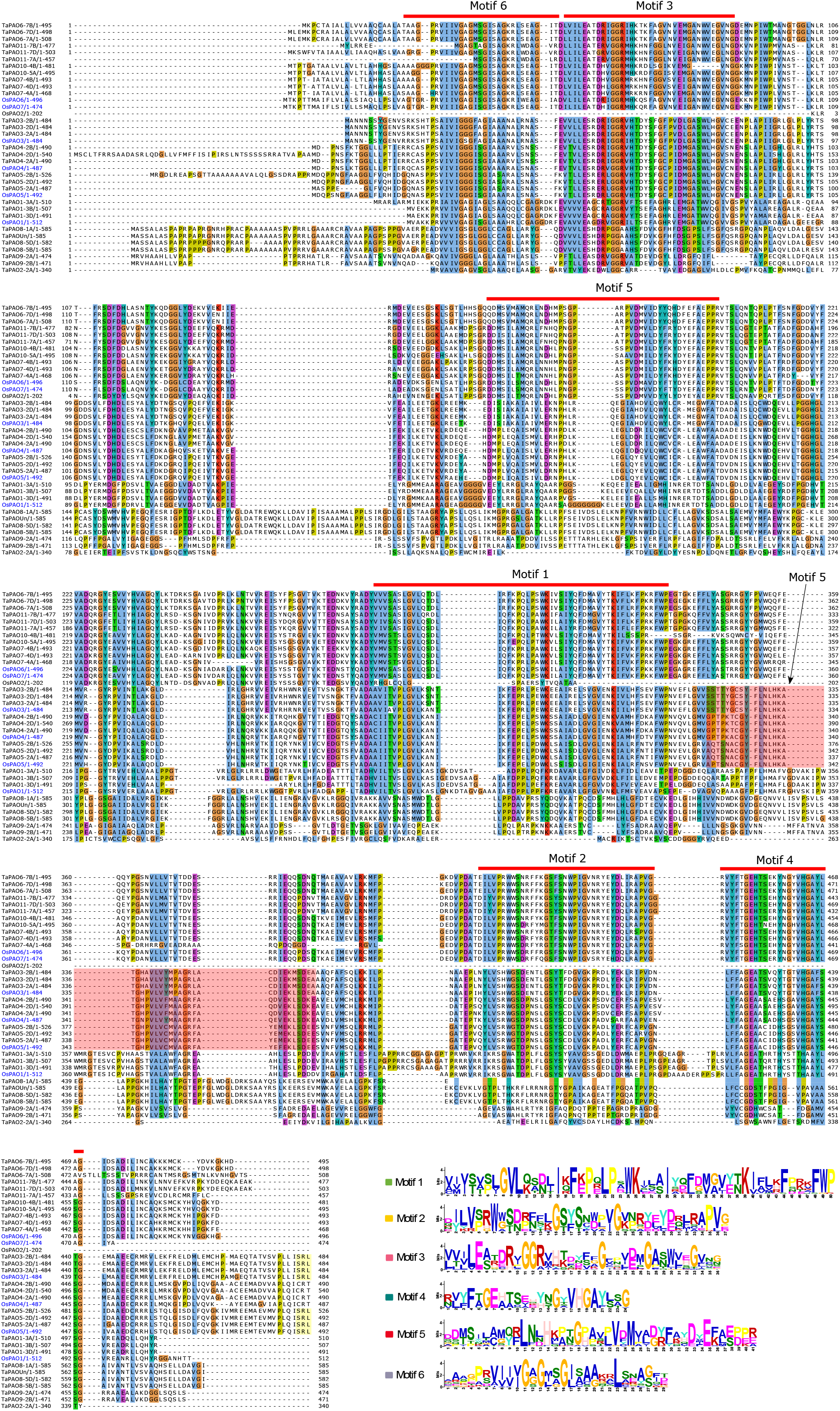
Multiple sequence alignment of wheat and rice PAO protein sequences. The locations and logos of the conserved domains of *TaPAO* genes identified by MEM are indicated. Searching in Pfam identified domains 1, 2, 3 and 4 as Flavin containing amine oxidoreductase; domain 6 as NAD_binding_8 and no result was found for domain 5.

### Expression profile analysis of *TaPAOs* under developmental stages

Analysis of expression profiles of *TaPAO* genes at various tissue and developmental stages using the expVIP data revealed that most *TaPAOs* are differentially expressed during developmental stages. For example, *TaPAO3-2A/2B*, TaPAO4-2A/2B/2D and *TaPAO5-2A/2B/2D* are highly expressed in specific tissues and developmental stages. The expression levels of *TaPAO11-7D* increased dramatically in some tissues such as leaf sheath, ligule, spike and spikelet during developmental stages. *TaPAO8-1A/5B/5D* genes also showed a clear tissue and developmental specific expression pattern and mainly downregulated in shoot, root and most parts of spike such as flower, ovary, anther, embryo and grain (Fig. 6). On the other hand, *TaPAO9-A/B/C*, *TaPAO7* and *TaPAO10* are less responsive to different conditions, tissues and developmental stages, although some homoeologs of these genes were active in some tissues and developmental stages (Fig. 6 and Fig. 7).

**Fig. 6.**
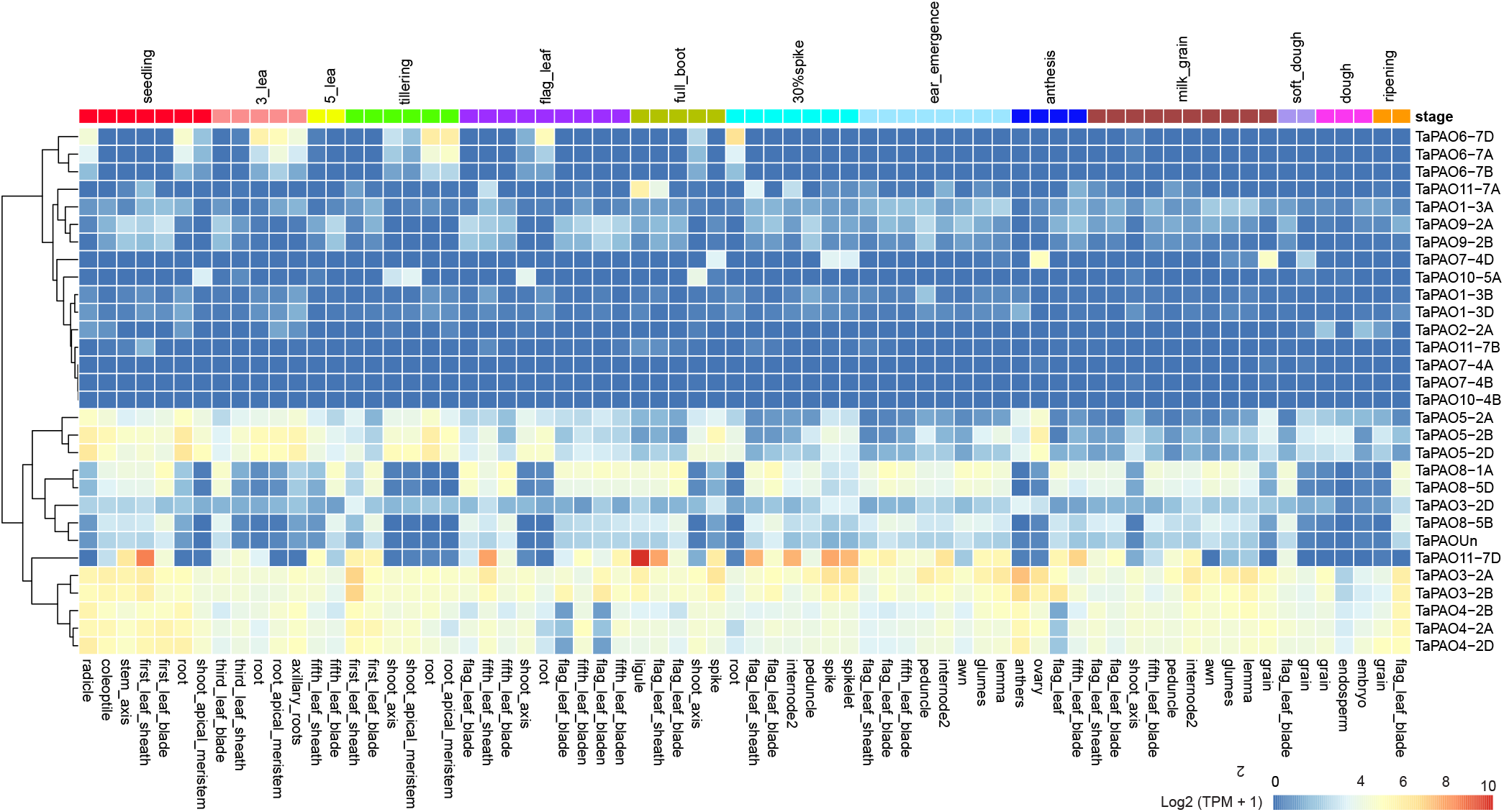
Log_2_ based expression levels for several *TaPAO* genes in different tissues during developmental stages. TPM values belong to Ramírez-González, Borrill (42) and retrieved from www.wheat-expression.com.

**Fig. 7.**
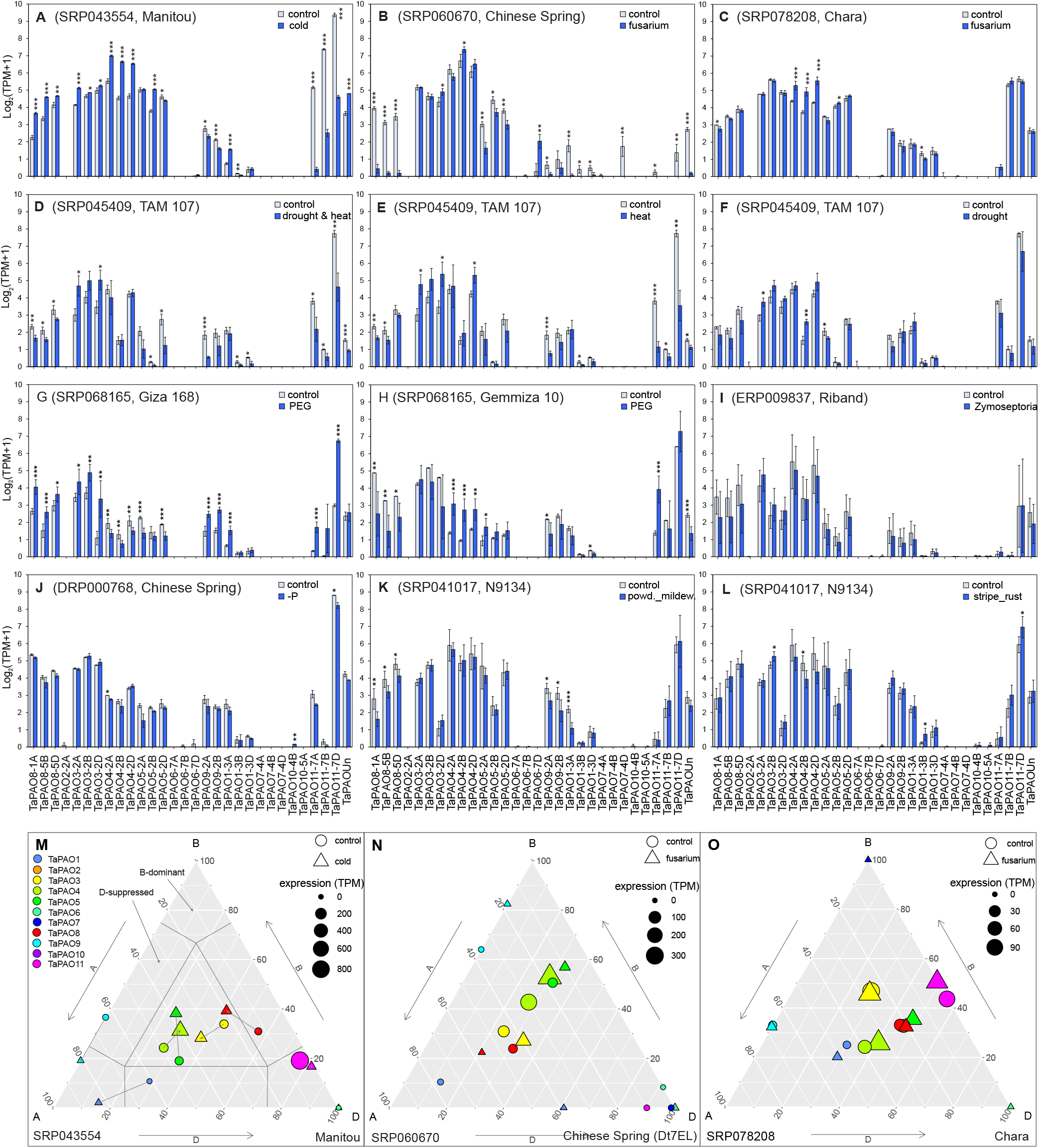
Barplots of the transcript expression rates (mean ± sd) of *TaPAO* genes in common wheat under different stress conditions including **A**) Leaf of ‘Manitou’ cultivar under normal (control) and cold stress conditions. **B**) ‘Chinese Spring’ cultivar 4 days after mock inoculation or inoculation with *F. graminearum*. **C**) Coleoptile-sheath-enclosed shoot tissue of common wheat ‘Chara’, 3 days after mock inoculation or inoculation with *F. graminearum*. **D**, **E** and **F**) seedlings of ‘TAM 107’ cultivar under a combined of heat and drought stress (40 °C and 20 % PEG-6000) and normal (22 °C) conditions (**D**) heat (40 °C) and normal (22 °C) conditions (**E**) and drought (20 % PEG-6000) and control (22 °C) conditions (**F**). **G** and **H**) Leaf tissue of ‘Ciza 168’ and ‘Gemmiza 10’ under control and PEG treatment conditions. **I**) Leaves of the “Riband” cultivar after mock inoculation (control) or inoculation with *Zymoseptoria tritici* isolate IPO323. **J**) Seedlings of the “Chinese Spring” cultivar 10 days after phosphorus starvation and under control conditions. **K**) Seedlings of the “N9134” cultivar 7 days after mock inoculation or after inoculation with powdery mildew. **L**) Seedlings of the “N9134” cultivar seven days after mock inoculation or inoculation with stripe rust. In each experiment ‘*’, ‘**’ and ‘***’ indicate statistically significant differences from control at 0.05, 0.01 and 0.005 significant levels, based on DESeq2 adjusted p-values. **M**, **N** and **O**) Ternary plot showing relative expression abundance of *TaPAO* genes under different stress conditions. In each ternary plot, a circle or a triangles reflects the relative contribution of homoeologs of a gene under the normal or stress condition respectively, and their sizes indicate the total expression in TPM. The data code for each study and the evaluated wheat cultivar are also indicated at the top (in barplots) or bottom (in ternary plots) of the subfigures.

### Expression profiles of TaPAOs under biotic and abiotic stresses

The differential expression of *TaPAOs* under biotic stresses (powdery mildew pathogen, *Zymoseptoria tritici*, stripe rust and *Fusarium graminearum* pathogen infections) and abiotic stresses (cold, heat, drought, heat and drought, phosphorus starvation and PEG) was assessed using the downloaded RNAseq data from expVIP. Results show that the expression of *TaPAO8, TaPAO3*, *TaPAO4*, *TaPAO5*, *TaPAO1-3A* and *TaPAOUn* was significantly upregulated in the leaf of the ‘Manitou’ cultivar under cold stress. However, *TaPAO11-7A/7B/7D* were downregulated under the same condition (Fig. 7A). Expression profiles of TaPAO-7D were also slightly downregulated under phosphorus starvation (Fig. 7J). Furthermore, the transcript expressions of *TaPAO3*, *TaPAO4* and *TaPAO5* homoeologs were significantly increased under heat or under a combination of heat and drought stresses relative to normal condition in seedling leaves of the ‘TAM 107’ cultivar (Fig. 7D and E), but these genes were not significantly affected by drought stress (Fig. 7F). An expression pattern relatively similar to heat stress was observed for *TaPAO3*, *TaPAO4* and *TaPAO5* homoeologs under PEG treatment, although they showed less expression abundance compared to under heat stress conditions (Fig. 7H and G). Contrary to the cold (A), heat and drought (B) and heat (C) stresses, the expression of *TaPAO11* homoeologs was significantly increased under PEG treatment, especially in the ‘Giza 168’ cultivar. Interestingly, *TaPAO3* and *TaPAO4* genes were differentially expressed between ‘Giza 168’ and ‘Gemmiza 10’: while the transcript levels of these genes decreased under PEG in ‘Giza 168’, expression of some genes, such as *TaPAO4* significantly increased under similar condition in ‘Gemmiza 10’.

Although some other genes nd homoeologs were differentially expressed in other experiments, high variation in the data prevented reliable conclusions (Fig. 7K, L). For example, the expression of *TaPAO4* homoeologs was significantly increased in coleoptile sheath enclosed shoot tissue of common wheat ‘Chara’ three days after inoculation with *F. graminearum* (Fig. 7C). Some *TaPAOs* were also differentially expressed between non-inoculated and inoculated leaves of the ‘N9134’ cultivar seven days after stripe rust and powdery mildew stress treatment (Fig. 7K, L).

Expression changes of *TaPAO* genes were also shown in ternary plots for the first three experiments of Fig. 7M, N and O. Ternary plots for the other *TaPAO* genes are presented in supplementary File 2, Fig. S1. Wheat ternary plots, provide an immediate view about the relative expression and abundance of homoeologous genes from each of the wheat three subgenomes. For example, the position of *TaPAO11* on the plot shows that it is dominantly expressed from the D subgenomes (supplementary file 2 Fig. S1, A-I and Fig. 7M), while *TaPAO1* is mainly expressed from the A subgenomes.

### Involvement of alternative splicing in *TaPAO* genes

To explore alternative splicing in *TaPAO* genes, the RNAseq data (45.31 Gb) from the leaves of common wheat cultivar ‘Manitou’ exposed to normal (23°C) and cold stress (4°C) conditions (accession number: SRP043554) was downloaded and aligned to the recent wheat reference genome. The overall alignment rate was 93.61%. Transcripts were assembled using StringTie. Differential transcript expression analysis and graphical displaying of alternative splice variants were done using the “Ballgown” package [54]. Compared to the number of splice variants mentioned for each gene in EnsemblPlants, novel isoforms were identified for 12 out of 30 *TaPAO* genes (Table 1). Because the wheat annotation file was used by StringTie during the assembly, most of the identified transcript should be due to alternative splicing. Structure and expression levels of distinct isoforms of the *TaPAO5-2D* gene under normal (23°C) and stress (4°C) conditions are illustrated in Fig. 8, where isoforms expressed at higher levels than the others are indicated by the darker color. Structure and expression levels of isoforms for the other *TaPAO* genes are presented in Supplementary File 3, Fig. S2. For most genes, different isoforms responded differently between normal and stress conditions (Fig. 8, supplementary file 3, Fig. S2). Among the *TaPAO* genes, we did not identify any isoforms that were available only in one condition.

**Fig. 8.**
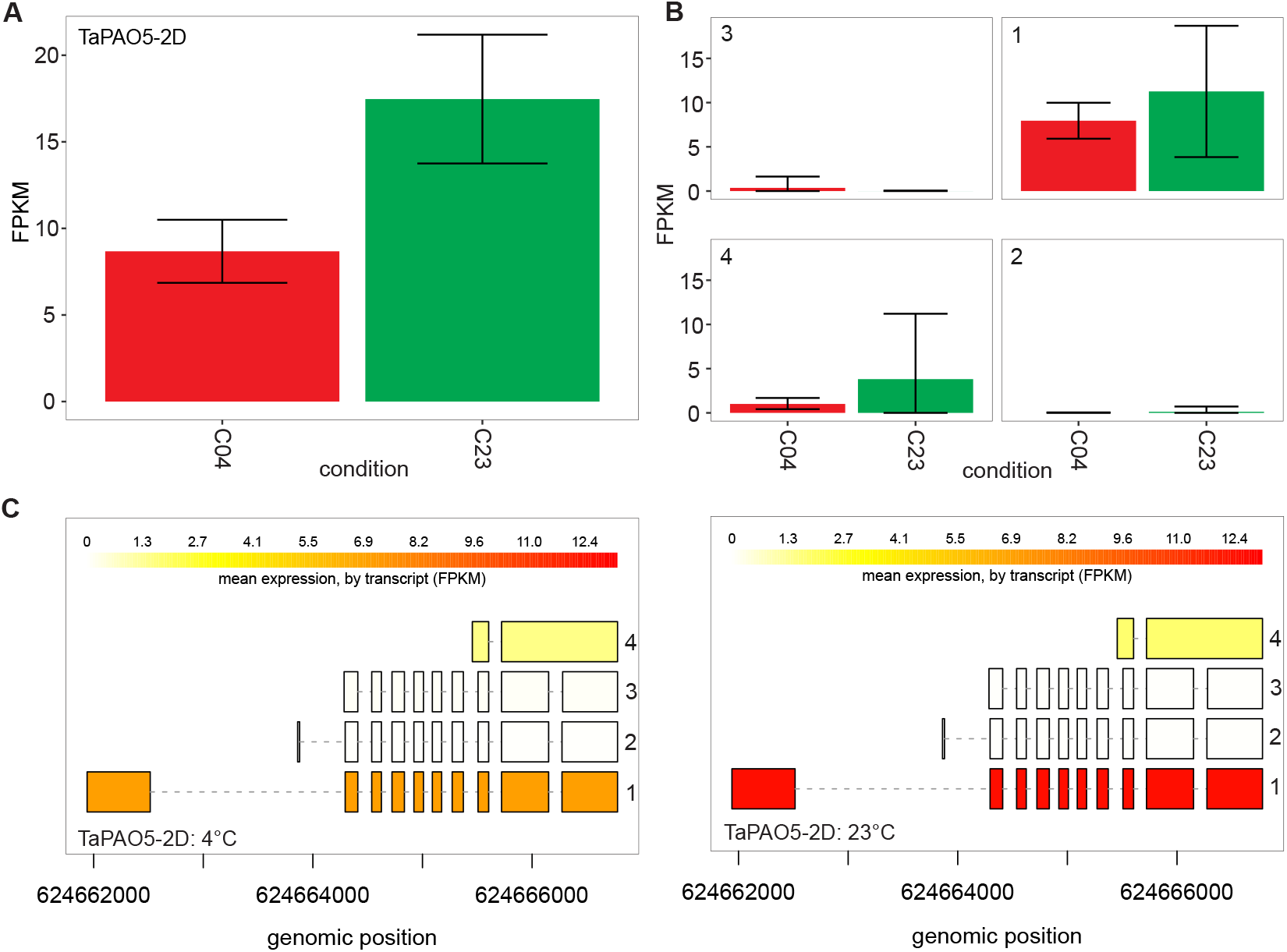
Expression levels in FPKM and the structure of distinct isoforms of the three *TaPAO5-2D* genes under normal (23°C) and stress (4°C) conditions from the SRP043554 experiment (A). Expression levels of isoforms are shown by barplots ± standard deviations (B) and in varying shades of yellow (C). Boxes represent exons and horizontal lines connecting exons represent introns.

## Discussion

### Structural characterization of polyamine oxidase genes (PAOs) in wheat

In the present study, we identified six *PAO* genes in diploid *T. urartu*, eight in diploid *Ae. tauschii* and 30 in hexaploid wheat (*T. aestivum*) by genome-wide approaches. We also structurally and functionally characterized the *TaPAO* genes using the publicly available RNAseq data. Previous studies have identified five PAO members in *A. thaliana* [55], seven in rice [4], two in barley [56], one in maize [57], seven in tomato [58], six in sweet orange [1], five in *Brachypodium distachyon* [59] and twelve in upland cotton [60]. AtPAO2~4, and OsPAO3~5, are believed to localize in peroxisomes based on possessing (S/A/C)(K/R/H)(L/M), in their C-termini which is a putative type-I peroxisomal targeting signal called PTS1 [4, 24]. Presence of SRL sequence in the C-termini of wheat TaPAO3 and TaPAO5 (Fig. 5) suggests that these proteins are localized in peroxisomes of wheat cells.

The identified *TaPAO* genes are distributed on 16 out of 21 wheat chromosomes plus the unassembled (Un) chromosome. As seen in the phylogenetic tree, each of the TaPAO homoeologous members aligned together in the same clade along with their *T. urartu* and *Ae. tauschii* orthologs (Fig. 2). The *TaPAO* genes generally showed an uneven distribution across the A, B, and D subgenomes. Similar biased distribution of gene family members is widespread. For example, *TaWD40*, *TaGST* and *TabZIP* family members are unevenly distributed across wheat chromosomes [61–63]. A high structural similarity of exon/intron structure between *TaPAO3*, *TaPAO4* and *TaPAO5*, and their close affinity at the distal end of the long arm of homoeologous group 2 suggest that a gene duplication event might be involved in the evolution of these genes [64].

### Expression profile analysis of *TaPAOs* during developmental stages

Tissue expression profile analysis revealed that many *TaPAOs* are expressed in a redundant manner in different tissues during developmental stages in bread wheat (Figure 5), supporting the idea that PAOs are involved in various tissues during all developmental processes in all living organisms [2, 6, 65].

### Expression profiles analysis of *TaPAOs* in response to abiotic stress

It is believed that PA molecules and PAOs also participate in responses to various abiotic stresses [6, 18, 65]. This has been specifically supported by the presence of putative *cis*-acting elements in the promoter region of polyamine biosynthetic genes including ADC and SAMDC which are regulated by transcription factors such as MYB, ABF and WRKY [66–68]. Concordantly, identification of consistently up- and downregulated expression patterns for a number of *TaPAOs* such as *TaPAO8, TaPAO4*, *TaPAO5* and *TaPAO11* under cold, drought or heat stresses suggest the involvement of *PAO* genes in multiple abiotic stress responses (Figure 6). Specifically, *TaPAOs* clearly responded to low and high temperatures. A similar temperature response has been suggested for *PAO* genes of cotton [60]. Similarly, *MdPAO2* expression was upregulated in apple fruit by elevating the CO_2_ concentrations under low-temperature/low-O_2_ storage [69]. In tomato, *SlPAOs* respond to abiotic stresses including heat, wounding, cold, drought, and salt [58].

In wheat, polyamine oxidases, were salt-induced in a salinity-tolerant genotype and showed higher expression compared with a salt-treated wild type, indicating that *TaPAOs* may play important roles in salinity tolerance as well [70]. TaPAOs have also been involved in osmotic stress: both abscisic acid pre-treatment and PEG induced osmotic stress, increased the Put, but decreased the Spm contents in wheat leaves, suggesting a connection between PA metabolism and abscisic acid signalling that leads to the controlled regulation and maintenance of Spd and Spm levels under osmotic stress in wheat seedlings [71]. Compared to high temperature alone, high temperature plus exogenous application of Spm and high temperature plus Spd significantly increased grain weight of a heat-resistant wheat variety by 19% and 5%, and of a heat-sensitive variety by 31% and 34%. Spm, Spd, and proline contents also increased significantly, while Put contents decreased during grain filling indicating that exogenous Spm and Spd could ameliorate heat damage during grain filling [72].

### Expression profile analysis of *TaPAOs* in response to biotic stress

Only a few *TaPAOs* significantly responded to biotic stresses during disease development but this was genotype and stress-type dependent and varied between experiments. This is not surprising because gene expression in response to biotic stress has been shown to vary significantly based on environmental conditions. For example, *F. graminearum* produces a different gene expression pattern when infecting diverse tissue types or at different stages of infection in wheat [73]. Differential gene expression patterns could also be dependent on the specific isolates infecting host genotypes [74].

Experiment SRP060670 (i.e. Fig. 6B) was the only case where *TaPAO11* genes which are located on the long arm of homoeologous group 7, were not expressed under both normal and *Fusarium* stress conditions. This result suggests that the examined wheat genotype in this case might be a ditelocentric addition line CS-7EL(7D) where the 7DL chromosome arm has been substituted by 7EL arm of *Thinipyrum elongatum* [43], subsequently affecting gene expression.

### Differential response of homoeologous genes

Differential response of homoeologous genes in allopolyploids is common when the plant is subjected to stresses. Here, unequal expression of homoeologs in response to stress was observed for some *TaPAO* genes such as *TaPAO11* under high temperature (Fig. 7D, E) and phorphorus starvation (Fig. 7J). Dong and Adams (2011) investigated the expression patterns of homoeologs in response to heat, cold, drought and high salt stresses in allotetraploid cotton (*Gossypium hirsutum*) and observed variation in the contribution of homoeologous genes to abiotic stresses [75]. Similarly, some homoeologs of *Coffea canephora* which are involved in the mannitol pathway, presented unequal contributions in response to drought, salt and heat stresses [76]. While PA-related genes play crucial roles in stress response, the mechanisms of this PA reaction are not clear. Some evidence suggests that PAO enzymes respond to stress mainly by modulating the homeostasis of reactive oxygen species (ROS) [1], but a clear understanding of the biochemical functions of PAO proteins requires more experimental investigation.

### Involvement of alternative splicing in *TaPAO* genes

Among the 30 *TaPAO* genes, 15 produced more than one isoform while only 3 *TaPAO* genes had alternative splice variants in EnsemblPlants. In total, 30 alternative splice variants were identified in wheat cultivar ‘Manitou’. Therefore, a major proportion of TaPAO transcript diversity is due to alternative splicing. Observation of a large fraction of novel isoforms in RNAseq data is common. It is believed that about 60% of intron-containing genes are alternatively spliced in plants [77, 78]. For example, 63% of intron containing genes are alternatively spliced in soybean, and on average, each AS gene contain six to seven AS events [78]. In common wheat, 200, 3576 and 4056 genes exhibited significant alternative splice pattern changes in response to drought, heat, and a combination of heat and drought stresses, respectively, implying that expression patterns of alternative splice variants are significantly altered by heat rather that drought [79]. Moreover, if RNAseq data from samples belonging to different developmental stages and extreme conditions were to be examined, a higher proportion of alternatively spliced genes and splice variants would likely be identified. Alternative splicing might also observed in different tissues and developmental stages [80]. But in the present study, all the *TaPAO* genes were constitutively alternatively spliced in all samples.

## Conclusion

We identified and characterized 30 *PAO* genes in common wheat that unevenly distributed across the wheat chromosomes. *TaPAO* genes were expressed redundantly in various tissues and developmental stages but a major fraction of *TaPAOs* responded significantly to abiotic stresses especially to temperature (i.e. heat and cold stresses). Some *TaPAOs* were also involved in responses to other stresses such as, powdery mildew, stripe rust and *Fusarium* infections in wheat. Overall, *TaPAOs* likely function in stress tolerances and play vital roles in different tissues and developmental stages. To understand the exact mechanisms of polyamine catabolism and biological functions of *TaPAOs*, more genetic and biochemical experiments are required. Our results provide a reference for further functional investigation of TaPAOs proteins.

## Acknowledgements

We thank Professor Annaliese Mason (Justus Liebig University, Germany) for providing helpful corrections. This work was supported by the University of Kurdistan.

## Author Contribution Statement

GM conceived and designed research. GM and FG conducted data analysis and wrote the manuscript.

---

**Supplementary File 1.**
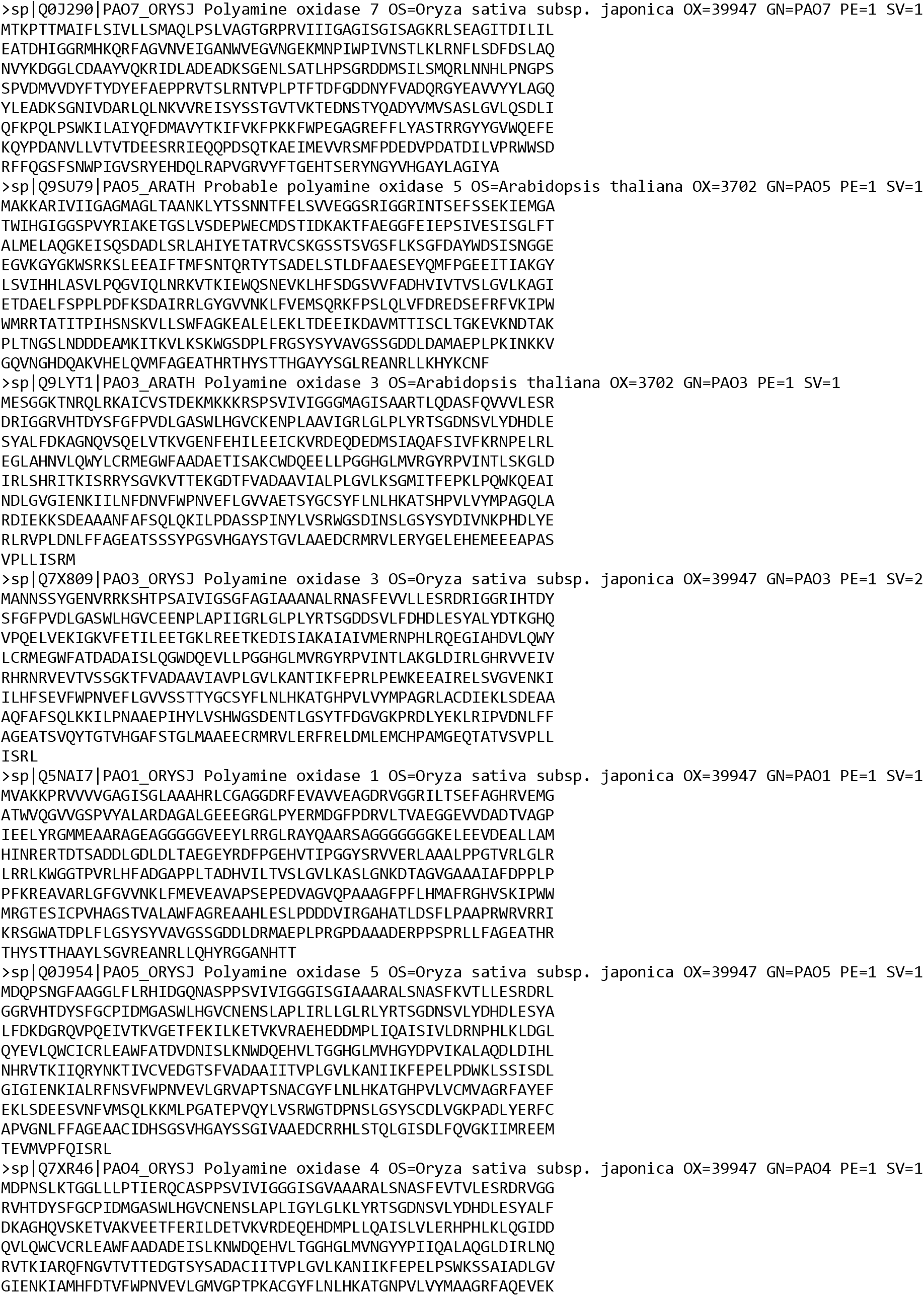
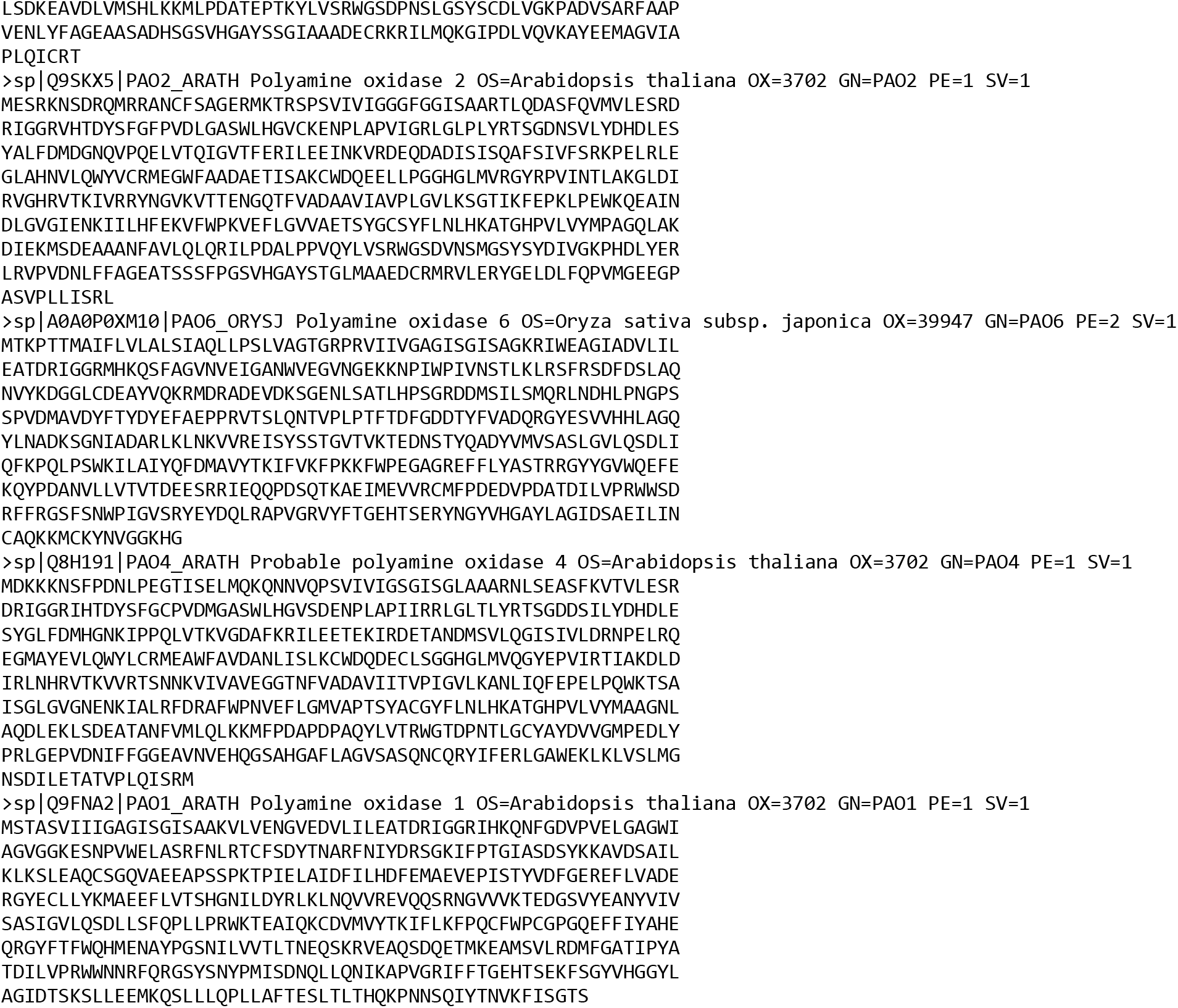

**Supplementary Fig. S1.**
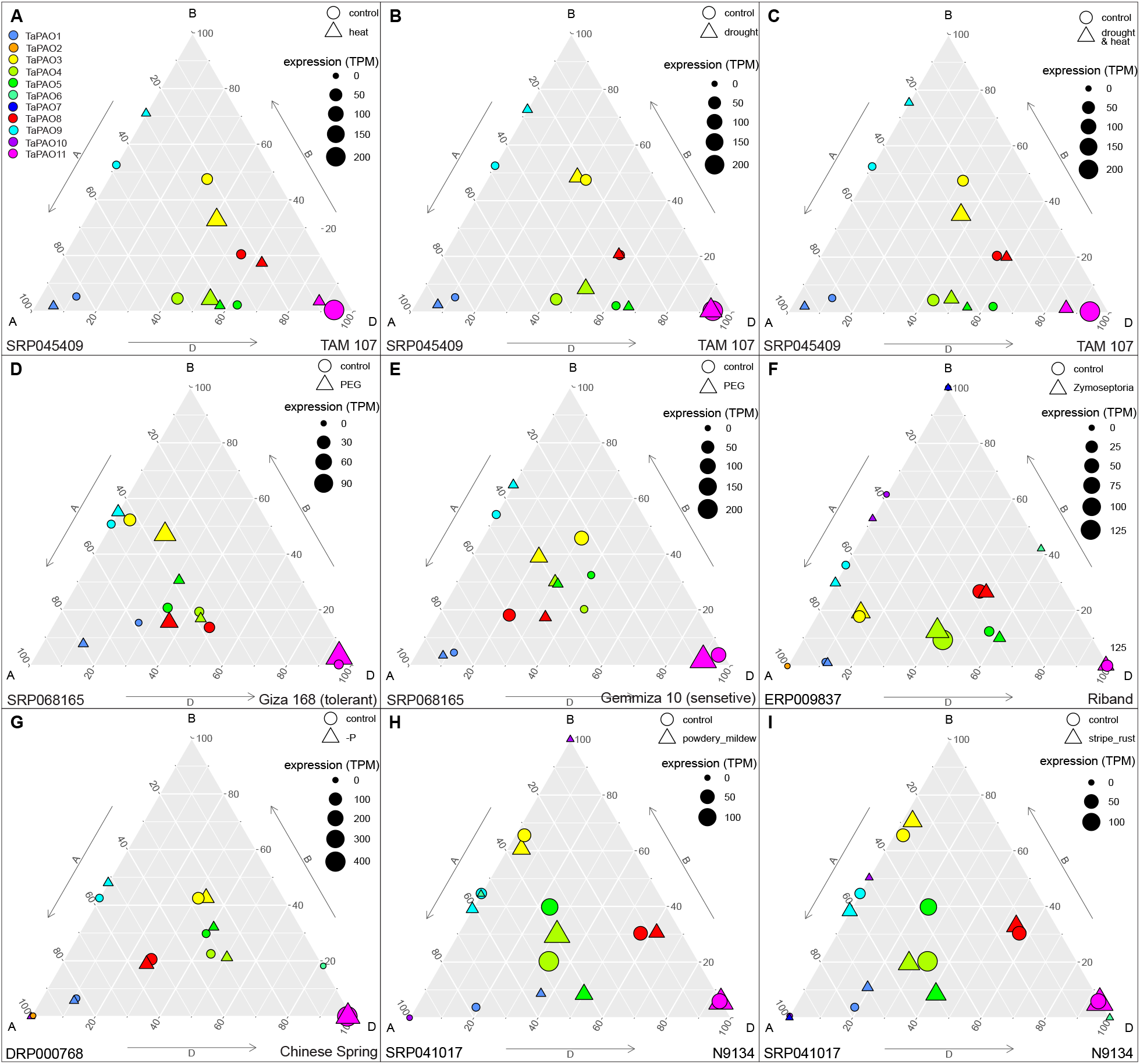
Ternary plot showing relative expression abundance of *TaPAO* genes under different stress conditions. In each ternary plot, a circle or a triangles reflects the relative contribution of homoeologs of a gene under normal or stress condition, respectively and their sizes indicate the total expression in TPM. The data code for each study and the evaluated wheat cultivar are also indicated at bottom of subfigures.

**Supplementary Fig. S2.**
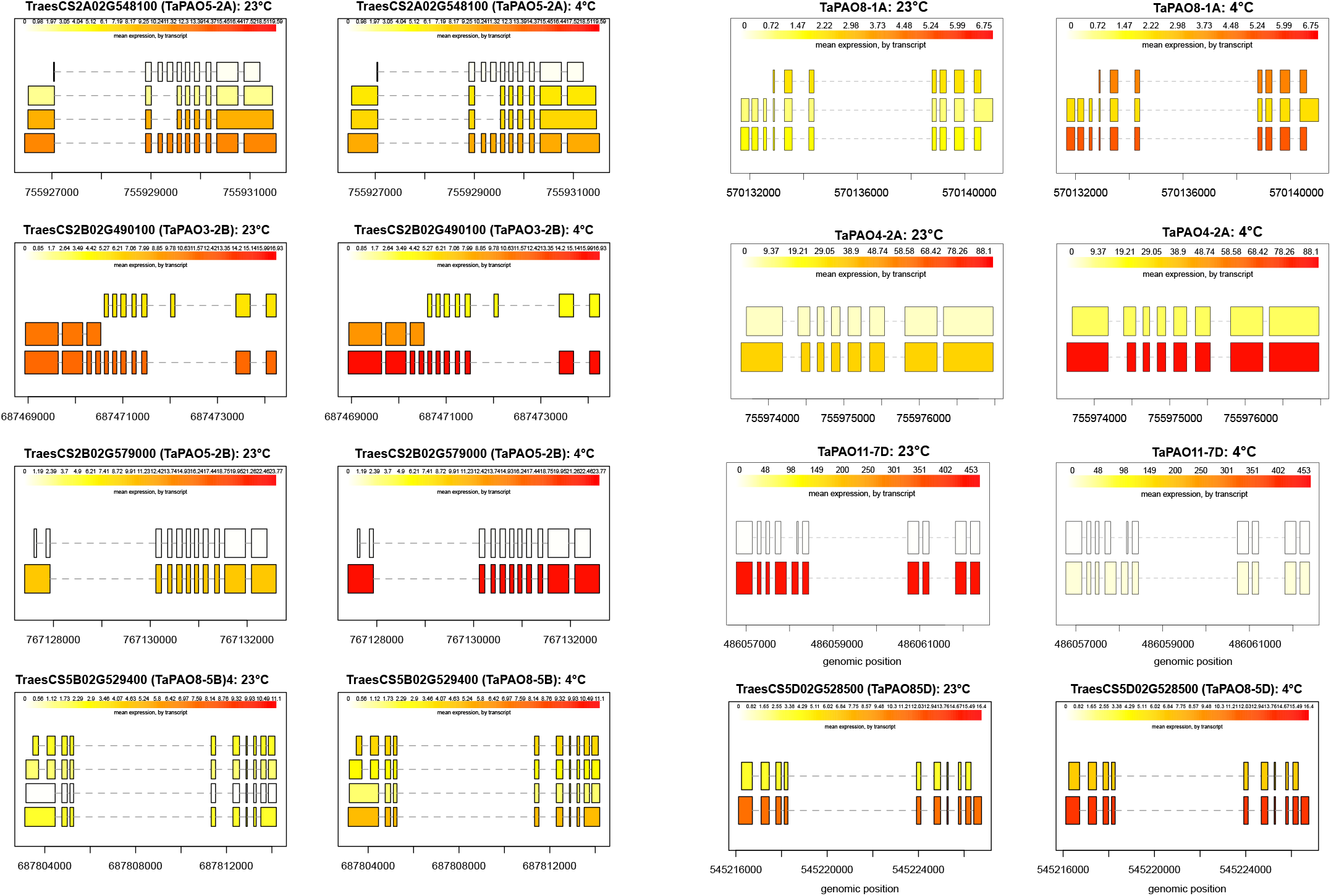
Structure and expression levels in FPKM of distinct isoforms of eight *TaPAO* genes in normal (23°C) and stress (4°C) from SRP043554 experiment. Expression levels are shown in varying shades of yellow.

## References

1. Liu J-H, Wang W, Wu H, Gong X, Moriguchi T. Polyamines function in stress tolerance: from synthesis to regulation. Frontiers in Plant Science. 2015;6:827.

2. Rangan P, Subramani R, Kumar R, Singh AK, Singh R. Recent advances in polyamine metabolism and abiotic stress tolerance. BioMed Research International. 2014;2014.

3. Corpas FJ, Del R ío LA, Palma JM. Plant peroxisomes at the crossroad of NO and H2O2 metabolism. Journal of integrative plant biology. 2019;61:803–16.

4. Ono Y, Kim DW, Watanabe K, Sasaki A, Niitsu M, Berberich T, et al. Constitutively and highly expressed *Oryza sativa* polyamine oxidases localize in peroxisomes and catalyze polyamine back conversion. Amino Acids. 2012;42:867–76.

5. Cona A, Rea G, Angelini R, Federico R, Tavladoraki P. Functions of amine oxidases in plant development and defence. Trends in Plant Science. 2006;11:80–8.

6. Alcázar R, Altabella T, Marco F, Bortolotti C, Reymond M, Koncz C, et al. Polyamines: molecules with regulatory functions in plant abiotic stress tolerance. Planta. 2010;231:1237–49.

7. Moschou P, Wu J, Cona A, Tavladoraki P, Angelini R, Roubelakis-Angelakis K. The polyamines and their catabolic products are significant players in the turnover of nitrogenous molecules in plants. Journal of Experimental Botany. 2012;63:5003–15.

8. Angelini R, Cona A, Federico R, Fincato P, Tavladoraki P, Tisi A. Plant amine oxidases “on the move”: an update. Plant Physiology and Biochemistry. 2010;48:560–4.

9. Mo H, Wang X, Zhang Y, Zhang G, Zhang J, Ma Z. Cotton polyamine oxidase is required for spermine and camalexin signalling in the defence response to *Verticillium dahliae*. The Plant Journal. 2015;83:962–75.

10. Sengupta A, Chakraborty M, Saha J, Gupta B, Gupta K. Polyamines: osmoprotectants in plant abiotic stress adaptation. Osmolytes and plants acclimation to changing environment: emerging omics technologies: Springer; 2016. p. 97–127.

11. Agudelo-Romero P, Bortolloti C, Pais MS, Tiburcio AF, Fortes AM. Study of polyamines during grape ripening indicate an important role of polyamine catabolism. Plant Physiology and Biochemistry. 2013;67:105–19.

12. Moschou PN, Paschalidis KA, Roubelakis-Angelakis KA. Plant polyamine catabolism: the state of the art. Plant Signaling and Behavior. 2008;3:1061–6.

13. Shelp BJ, Deyman KL, DeEll JR, Bozzo GG. Polyamine homeostasis in apple fruit stored under multiple abiotic stresses. Canadian Journal of Plant Science. 2018;99:88–92.

14. Cui J, Pottosin I, Lamade E, Tcherkez G. What is the role of putrescine accumulated under potassium deficiency? Plant, Cell and Environment. 2020:1–17. doi: 10.1111/pce.13740.

15. Sequera-Mutiozabal M, Tiburcio AF, Alcázar R. Drought Stress Tolerance in Relation to Polyamine Metabolism in Plants. In: Hossain MA, Wani SH, Bhattacharjee S, Burritt DJ, Tran L-SP, editors. Drought Stress Tolerance in Plants, Vol 1: Physiology and Biochemistry. Cham: Springer International Publishing; 2016. p. 267–86.

16. Fu X-Z, Chen C-W, Wang Y, Liu J-H, Moriguchi T. Ectopic expression of MdSPDS1 in sweet orange (*Citrus sinensis* Osbeck) reduces canker susceptibility: involvement of H2 O2 production and transcriptional alteration. BMC Plant Biology. 2011;11:55.

17. Mitsuya Y, Takahashi Y, Berberich T, Miyazaki A, Matsumura H, Takahashi H, et al. Spermine signaling plays a significant role in the defense response of *Arabidopsis thaliana* to cucumber mosaic virus. Journal of Plant Physiology. 2009;166:626–43.

18. Yu Y, Zhou W, Zhou K, Liu W, Liang X, Chen Y, et al. Polyamines modulate aluminum-induced oxidative stress differently by inducing or reducing H2O2 production in wheat. Chemosphere. 2018;212:645–53.

19. Ozawa R, Bertea CM, Foti M, Narayana R, Arimura G-I, Muroi A, et al. Exogenous polyamines elicit herbivore-induced volatiles in lima bean leaves: involvement of calcium, H2O2 and Jasmonic acid. Plant and Cell Physiology. 2009;50:2183–99.

20. Hatmi S, Trotel-Aziz P, Villaume S, Couderchet M, Clément C, Aziz A. Osmotic stress-induced polyamine oxidation mediates defence responses and reduces stress-enhanced grapevine susceptibility to *Botrytis cinerea*. Journal of Experimental Botany. 2014;65:75–88.

21. Xu X, Shi G, Jia R. Changes of polyamine levels in roots of *Sagittaria sagittifolia* L. under copper stress. Environmental Science and Pollution Research. 2012;19:2973–82.

22. Yang H, Shi G, Wang H, Xu Q. Involvement of polyamines in adaptation of *Potamogeton crispus* L. to cadmium stress. Aquatic Toxicology. 2010;100:282–8.

23. Cervelli M, Caro OD, Penta AD, Angelini R, Federico R, Vitale A, et al. A novel C‐terminal sequence from barley polyamine oxidase is a vacuolar sorting signal. The Plant Journal. 2004;40:410–8.

24. Moschou PN, Sanmartin M, Andriopoulou AH, Rojo E, Sanchez-Serrano JJ, Roubelakis-Angelakis KA. Bridging the gap between plant and mammalian polyamine catabolism: a novel peroxisomal polyamine oxidase responsible for a full back-conversion pathway in *Arabidopsis*. Plant Physiology. 2008;147:1845–57.

25. Kim DW, Watanabe K, Murayama C, Izawa S, Niitsu M, Michael AJ, et al. Polyamine oxidase5 regulates *Arabidopsis* growth through thermospermine oxidase activity. Plant Physiology. 2014;165:1575–90.

26. Ahou A, Martignago D, Alabdallah O, Tavazza R, Stano P, Macone A, et al. A plant spermine oxidase/dehydrogenase regulated by the proteasome and polyamines. Journal of Experimental Botany. 2014;65:1585–603.

27. Fincato P, Moschou PN, Ahou A, Angelini R, Roubelakis-Angelakis KA, Federico R, et al. The members of Arabidopsis thalianaPAO gene family exhibit distinct tissue-and organ-specific expression pattern during seedling growth and flower development. Amino Acids. 2012;42:831–41.

28. Takahashi T, Kakehi J-I. Polyamines: ubiquitous polycations with unique roles in growth and stress responses. Annals of Botany. 2010;105:1–6.

29. Finn RD, Clements J, Arndt W, Miller BL, Wheeler TJ, Schreiber F, et al. HMMER web server: 2015 update. Nucleic Acids Research. 2015;43:W30–W8.

30. Spedaletti V, Polticelli F, Capodaglio V, Schininà ME, Stano P, Federico R, et al. Characterization of a lysine-specific histone demethylase from *Arabidopsis thaliana*. Biochemistry. 2008;47:4936–47.

31. Yu Y, Ouyang Y, Yao W. shinyCircos: an R/Shiny application for interactive creation of Circos plot. Bioinformatics 2017;34:1229–31.

32. Gasteiger E, Hoogland C, Gattiker A, Duvaud Se, Wilkins MR, Appel RD, et al. Protein Identification and Analysis Tools on the ExPASy Server. In: Walker JM, editor. The Proteomics Protocols Handbook. Totowa, NJ: Humana Press; 2005. p. 571–607.

33. Hu B, Jin J, Guo A, Zhang H, Luo J, Gao G. GSDS 2.0: An upgraded gene feature visualization server. Bioinformatics. 2015;31:1296–7.

34. El-Gebali S, Mistry J, Bateman A, Eddy SR, Luciani A, Potter SC, et al. The Pfam protein families database in 2019. Nucleic Acids Research. 2019;47:D427–D32.

35. Bailey TL, Boden M, Buske FA, Frith M, Grant CE, Clementi L, et al. MEME SUITE: tools for motif discovery and searching. Nucleic Acids Research. 2009;37:W202–W8.

36. Chen C, Chen H, He Y, Xia R. TBtools, a Toolkit for Biologists integrating various biological data handling tools with a user-friendly interface. bioRxiv 2018:289660.

37. Waterhouse AM, Procter JB, Martin DM, Clamp M, Barton G. Jalview Version 2—a multiple sequence alignment editor and analysis workbench. Bioinformatics. 2009;25(9):1189–91.

38. Bodenhofer U, Bonatesta E, Horejš-Kainrath C, Hochreiter S. msa: an R package for multiple sequence alignment. Bioinformatics. 2015;31(24):3997–9.

39. Paradis E, Schliep K. ape 5.0: an environment for modern phylogenetics and evolutionary analyses in R. Bioinformatics. 2019;35(3):526–8.

40. Yu G, Smith DK, Zhu H, Guan Y, Lam TTY, Evolution. ggtree: an R package for visualization and annotation of phylogenetic trees with their covariates and other associated data. Methods in Ecology. 2017;8:28–36.

41. Borrill P, Ramirez-Gonzalez R, Uauy C. expVIP: a customizable RNA-seq data analysis and visualization platform. Plant Physiology. 2016;170:2172–86.

42. Ramírez-González RH, Borrill P, Lang D, Harrington SA, Brinton J, Venturini L, et al. The transcriptional landscape of polyploid wheat. Science. 2018;361(6403):eaar6089.

43. Gou L, Hattori J, Fedak G, Balcerzak M, Sharpe A, Visendi P, et al. Development and validation of *Thinopyrum elongatum*–expressed molecular markers specific for the long arm of chromosome 7E. Crop Science. 2016;56:354–64.

44. Powell JJ, Fitzgerald TL, Stiller J, Berkman PJ, Gardiner DM, Manners JM, et al. The defence-associated transcriptome of hexaploid wheat displays homoeolog expression and induction bias. Plant biotechnology journal. 2017;15:533–43. Epub 2016/10/14. doi: 10.1111/pbi.12651. PubMed PMID: 27735125; PubMed Central PMCID: PMCPMC5362679.

45. Li Q, Zheng Q, Shen W, Cram D, Fowler DB, Wei Y, et al. Understanding the biochemical basis of temperature-induced lipid Pathway adjustments in plants. The Plant Cell. 2015;27:86–103.

46. Rudd JJ, Kanyuka K, Hassani-Pak K, Derbyshire M, Andongabo A, Devonshire J, et al. Transcriptome and metabolite profiling of the infection cycle of *Zymoseptoria tritici* on wheat reveals a biphasic interaction with plant immunity involving differential pathogen chromosomal contributions and a variation on the hemibiotrophic lifestyle definition. Plant Physiology. 2015;167:1158–85. doi: 10.1104/pp.114.255927 %J Plant Physiology.

47. Liu Z, Xin M, Qin J, Peng H, Ni Z, Yao Y, et al. Temporal transcriptome profiling reveals expression partitioning of homeologous genes contributing to heat and drought acclimation in wheat (*Triticum aestivum* L.). BMC Plant Biology. 2015;15:152.

48. Oono Y, Kobayashi F, Kawahara Y, Yazawa T, Handa H, Itoh T, et al. Characterisation of the wheat (*Triticum aestivum* L.) transcriptome by de novo assembly for the discovery of phosphate starvation-responsive genes: gene expression in Pi-stressed wheat. BMC Genomics. 2013;14:77. doi: 10.1186/1471-2164-14-77.

49. Zhang H, Yang Y, Wang C, Liu M, Li H, Fu Y, et al. Large-scale transcriptome comparison reveals distinct gene activations in wheat responding to stripe rust and powdery mildew. BMC Genomics. 2014;15:898.

50. Love MI, Huber W, Anders S. Moderated estimation of fold change and dispersion for RNA-seq data with DESeq2. Genome Biology. 2014;15:550.

51. Hamilton NE, Ferry M. ggtern: ternary diagrams using ggplot2. Journal of Statistical Software. 2018;87:1–17.

52. Appels R, Eversole K, Feuillet C, Keller B, Rogers J, Stein N, et al. Shifting the limits in wheat research and breeding using a fully annotated reference genome. Science. 2018;361:eaar7191.

53. Pertea M, Kim D, Pertea GM, Leek JT, Salzberg SL. Transcript-level expression analysis of RNA-seq experiments with HISAT, StringTie and Ballgown. Nature Protocols. 2016;11:1650

54. Frazee AC, Pertea G, Jaffe AE, Langmead B, Salzberg SL, Leek JT. Ballgown bridges the gap between transcriptome assembly and expression analysis. Nature Biotechnology. 2015;33:243.

55. Fincato P, Moschou PN, Spedaletti V, Tavazza R, Angelini R, Federico R, et al. Functional diversity inside the *Arabidopsis* polyamine oxidase gene family. Journal of Experimental Botany. 2011;62:1155–68.

56. Cervelli M, Cona A, Angelini R, Polticelli F, Federico R, Mariottini P. A barley polyamine oxidase isoform with distinct structural features and subcellular localization. European Journal of Biochemistry. 2001;268:3816–30.

57. Cervelli M, Tavladoraki P, Di Agostino S, Angelini R, Federico R, Mariottini P. Isolation and characterization of three polyamine oxidase genes from *Zea mays*. Plant Physiology and Biochemistry. 2000;38:667–77.

58. Hao Y, Huang B, Jia D, Mann T, Jiang X, Qiu Y, et al. Identification of seven polyamine oxidase genes in tomato (*Solanum lycopersicum* L.) and their expression profiles under physiological and various stress conditions. Journal of Plant Physiology. 2018;228:1–11.

59. Takahashi Y, Ono K, Akamine Y, Asano T, Ezaki M, Mouri I. Highly-expressed polyamine oxidases catalyze polyamine back conversion in *Brachypodium distachyon*. Journal of Plant Research. 2018;131:341–8.

60. Cheng X-Q, Zhu X-F, Tian W-G, Cheng W-H, Sun J, Jin S-X, et al. Genome-wide identification and expression analysis of polyamine oxidase genes in upland cotton (*Gossypium hirsutum* L.). Plant Cell, Tissue and Organ Culture. 2017;129:237–49.

61. Hu R, Xiao J, Gu T, Yu X, Zhang Y, Chang J, et al. Genome-wide identification and analysis of WD40 proteins in wheat (*Triticum aestivum* L.). BMC Genomics. 2018;19:803.

62. Wang R, Ma J, Zhang Q, Wu C, Zhao H, Wu Y, et al. Genome-wide identification and expression profiling of glutathione transferase gene family under multiple stresses and hormone treatments in wheat (*Triticum aestivum* L.). BMC Genomics. 2019;20:1–15.

63. Li X, Gao S, Tang Y, Li L, Zhang F, Feng B, et al. Genome-wide identification and evolutionary analyses of bZIP transcription factors in wheat and its relatives and expression profiles of anther development related TabZIP genes. BMC Genomics. 2015;16:976.

64. Huo N, Zhang S, Zhu T, Dong L, Wang Y, Mohr T, et al. Gene duplication and evolution dynamics in the homeologous regions harboring multiple prolamin and resistance gene families in hexaploid wheat. Frontiers in Plant Science. 2018;9:673.

65. Yu Z, Jia D, Liu T. Polyamine oxidases play various roles in plant development and abiotic stress tolerance. Plants. 2019;8:184.

66. Basu S, Roychoudhury A, Sengupta DN. Identification of trans-acting factors regulating SamDC expression in *Oryza sativa*. Biochemical and Biophysical Research Communications. 2014;445:398–403.

67. Gong X, Zhang J, Hu J, Wang W, Wu H, Zhang Q, et al. FcWRKY 70, a WRKY protein of *Fortunella crassifolia*, functions in drought tolerance and modulates putrescine synthesis by regulating arginine decarboxylase gene. Plant, Cell and Environment. 2015;38:2248–62.

68. Sun P, Zhu X, Huang X, Liu J-H. Overexpression of a stress-responsive MYB transcription factor of *Poncirus trifoliata* confers enhanced dehydration tolerance and increases polyamine biosynthesis. Plant Physiology and Biochemistry. 2014;78:71–9.

69. Brikis CJ, Zarei A, Chiu GZ, Deyman KL, Liu J, Trobacher CP, et al. Targeted quantitative profiling of metabolites and gene transcripts associated with 4-aminobutyrate (GABA) in apple fruit stored under multiple abiotic stresses. Horticulture Research. 2018;5:1–14.

70. Xiong H, Guo H, Xie Y, Zhao L, Gu J, Zhao S, et al. RNAseq analysis reveals pathways and candidate genes associated with salinity tolerance in a spaceflight-induced wheat mutant. Scientific Reports. 2017;7(1):1–13.

71. Pál M, Tajti J, Szalai G, Peeva V, Végh B, Janda T. Interaction of polyamines, abscisic acid and proline under osmotic stress in the leaves of wheat plants. Scientific Reports. 2018;8(1):12839. doi: 10.1038/s41598-018-31297-6.

72. Jing J, Guo S, Li Y, Li W. The alleviating effect of exogenous polyamines on heat stress susceptibility of different heat resistant wheat (*Triticum aestivum* L.) varieties. Scientific Reports. 2020;10(1):7467.

73. Zhang X-W, Jia L-J, Zhang Y, Jiang G, Li X, Zhang D, et al. In planta stage-specific fungal gene profiling elucidates the molecular strategies of *Fusarium graminearum* growing inside wheat coleoptiles. The Plant Cell. 2012;24:5159–76.

74. Hofstad AN, Nussbaumer T, Akhunov E, Shin S, Kugler KG, Kistler HC, et al. Examining the transcriptional response in wheat Fhb1 near-isogenic lines to *Fusarium graminearum* infection and deoxynivalenol treatment. The Plant Genome. 2016;9:10.3835.

75. Dong S, Adams KL. Differential contributions to the transcriptome of duplicated genes in response to abiotic stresses in natural and synthetic polyploids. New Phytologist. 2011;190:1045–57.

76. de Carvalho K, Petkowicz CL, Nagashima GT, Bespalhok Filho JC, Vieira LG, Pereira LF, et al. Homeologous genes involved in mannitol synthesis reveal unequal contributions in response to abiotic stress in *Coffea arabica*. Molecular Genetics and Genomics. 2014;289:951–63.

77. Reddy AS, Marquez Y, Kalyna M, Barta A. Complexity of the alternative splicing landscape in plants. . Plant Cell. 2013;25:3657–83.

78. Shen Y, Zhou Z, Wang Z, Li W, Fang C, Wu M, et al. Global dissection of alternative splicing in paleopolyploid soybean. The Plant Cell. 2014;26:996–1008.

79. Liu Z, Qin J, Tian X, Xu S, Wang Y, Li H, et al. Global profiling of alternative splicing landscape responsive to drought, heat and their combination in wheat (*Triticum aestivum* L.). Plant biotechnology journal. 2018;16:714–26. Epub 2017/08/24. PubMed PMID: 28834352; PubMed Central PMCID: PMCPMC5814593.

80. Yoshimura K, Yabuta Y, Ishikawa T, Shigeoka S. Identification of a cis element for tissue-specific alternative splicing of chloroplast ascorbate peroxidase pre-mRNA in higher plants. Journal of Biological Chemistry. 2002;277:40623–32.

